# Multiomic analisys reveals that viroid infection induces a temporal reprograming of plant-defence mechanisms at multiple regulatory levels

**DOI:** 10.1101/2022.01.06.475203

**Authors:** Joan Márquez-Molins, Pascual Villalba-Bermell, Julia Corell-Sierra, Vicente Pallás, Gustavo Gómez

**Affiliations:** Institute for Integrative Systems Biology (I2SysBio), Consejo Superior de Investigaciones Científicas (CSIC) - Universitat de València (UV), Parc Científic, Cat. Agustín Escardino 9, 46980 Paterna, Spain; Instituto de Biología Molecular y Celular de Plantas (IBMCP), Consejo Superior de Investigaciones Científicas (CSIC) - Universitat Politècnica de València, CPI 8E, Av. de los Naranjos s/n, 46022 Valencia, Spain

**Keywords:** Viroid-host interactions, epigenetic epidemiology, host transcriptional regulation in response to pathogens, global response to biotic stress, systems biology and diseases

## Abstract

Viroids are circular RNAs of minimal complexity compelled to subvert plant-regulatory networks to accomplish their infectious process. Studies focused on the response to viroid infection have mostly addressed specific regulatory levels and considered a unique infection time. Thus, much remains to be done to understand the temporal evolution and complex nature of viroid-host interactions. Here we present an integrative analysis of the temporal evolution and intensity of the genome-wide alterations in cucumber plants infected with hop stunt viroid (HSVd) by integrating differential host transcriptome, sRNAnome and methylome. Our results support that HSVd promotes the redesign of the cucumber regulatory-pathways predominantly affecting specific regulatory layers at different infection-phases. The initial response was characterized by a reconfiguration of the host-transcriptome by differential exon usage, followed by a progressive transcriptional down-regulation modulated by epigenetic changes. Regarding endogenous small RNAs, the alterations were limited and mainly occur at the late stage. The most significant host-alterations were predominantly related to the down-regulation of transcripts involved in plant-defence mechanisms, the restriction of pathogen-movement and the systemic spreading of defence signals. Altogether our data evidence the existence of a dynamic and yet poorly known arms race between the host and the viroid. We expect that these data constituting the first comprehensive map of the plant responses to a viroid infection contribute to elucidate the molecular basis of this multifaceted defence and counter-defence layout.

## INTRODUCTION

Viroids (the smallest infectious agents known to date) comprise an intriguing group of plant pathogens affecting a varied range of hosts worldwide. These sub-viral pathogens are classified into two families, *Pospiviroidae* whose replication takes place in the nucleus, and the *Avsunviroidae*, that replicates in the chloroplast [1]. As naked circular viroids are compelled to closely interact with diverse (many yet unknown) host factors in order to fulfil their infectious cycle in the infected plants [2–4]. However, how these pathogenic RNAs alter the host development and physiology inducing disease symptoms is still an unsolved question [5–7].

In general, plants respond to stress conditions (including pathogen infection) through a complex reprogramming of their transcriptional activity. Gene regulation is a complex process modulated by a series of coordinated events involving multiple control layers such as: epigenetic modifications, modulation of the transcriptional activity and small RNA interference [8–10]. These different regulatory levels have been categorized as stress-responsive structures termed as Environmental Gene Regulatory Influence Networks (EGRINs) [11].

Increasing evidence indicate that viroids can subvert different host-EGRINs and consequently promote the emergence of the phenotypic alterations recognized (in certain viroid-host interactions) as symptoms. In this sense, diverse studies have provided support to the notion that during the infectious process, viroids promote significant alterations in diverse plant regulatory pathways [5,7].

Although it has been proposed that viroids (presumably because of the highly compacted and structured genomic circular RNA), are resistant to degradation mediated by RNA-silencing [12–14], early reports evidenced that viroid-derived sRNAs (vd-sRNAs) (probably arising from the DCL-mediated processing of replication intermediates), are detected in plants infected by members of both nuclear and chloroplastic families [15–17]. The possibility that the vd-sRNAs could guide the silencing of plant-endogenous transcripts triggering the induction of disease symptoms, was then proposed as a plausible hypothesis to explain the basis of viroid-induced pathogenesis [17–19]. Several research groups have demonstrated in the last years the involvement of vd-sRNAs in the down-regulation of host transcripts [20–28], thus emphasizing the role of RNA silencing in viroid-induced symptoms. In addition, it has been recently proposed that viroid infection might trigger the production of secondary siRNAs able to recognize host transcripts that are not direct targets of the vd-sRNAs [21]. Also related to viroid-pathogenesis and RNA-silencing, alterations in the accumulation of certain endogenous miRNAs have been reported in diverse viroid-host interactions [27,29–35].

Modulation (in both positive and/or negative sense) of the transcriptional activity constitutes the core of the plant response to abiotic and biotic stress conditions [36,37]. Changes at the transcriptomic level have been described in several plants infected by viroids of both *Pospiviroidae* and *Avsunviroidae* families [38–44]. Altogether, these studies revealed a global alteration in the expression of host genes related to defense, response to stress, hormone homeostasis and signaling, biosynthesis of cell wall compounds, and RNA metabolism, among others [5].

A growing body of evidence points toward the existence of a close interplay between infection and epigenetic alterations in the host (plant or animal) genome [10,45]. Consequently, epigenetic epidemiology has emerged as a promising area for future research on infectious diseases [46]. Recent studies have evidenced dynamic changes in host DNA methylation patterns (mainly rDNA and transposable elements) upon viroid infection in both somatic [47,48] and reproductive [49] tissues. The changes of the host-genome epigenetic regulation have been connected (in infected cucumber plants) with viroid recruiting and functionally subverting the host HISTONE DEACETYLASE 6 (HDA6) [48]. Alteration of the host plant methylome has been also described to occur in response to infection by viruses [50] and bacteria [51], supporting that regulation of host DNA methylation may be part of an evolutionary conserved immune response in plants [52].

The exposed above evidenced that the advent of the high throughput technologies has favored the study of the viroid-host interactions at a molecular level. However, such experimental approaches have been generally centered on the analysis of the viroid-induced changes at a single host-regulatory level and/or considering a particular infection time (mainly correlated to advanced disease stages). Therefore, our knowledge about the functional diversity and temporal dynamics of the global plant response to viroid infection is scarce [3].

Here we have performed an integrative analysis of the timing and intensity of the genome-wide alterations in cucumber (*Cucumis sativus*) plants in response to hop stunt viroid (HSVd) infection [53]. Our study has been focused on the temporal dynamics (at three infection time-points) of the plant response to viroid infection at three different regulatory levels of the gene expression network: *i)* small RNA interference, *ii)* modulation of the transcriptional activity and *iii)* epigenetic modifications. The obtained results evidenced that HSVd infection promotes a dynamic redesign of the cucumber regulatory pathways that affects these regulatory layers distinctively depending on the infection phase.

## RESULTS

### Viroid infection

To uncover the dynamic of the host regulatory response to HSVd in cucumber we analyzed apical leaves of infected plants at 10-, 17- and 24-days post inoculation (dpi). Mock-inoculated plants were used as control. Time points were arbitrarily selected as representative of the early, intermediate and late HSVd infection phases, respectively. As expected, viroid induced symptoms were evident in HSVd-infected plants at 24 dpi (Figure 1A). A total of 54 libraries (18 for each sRNAs, long transcripts and methyl-C analysis) consisting of three bio replicates per time point of infected and control plants were obtained (Supplementary Figure 1). The information about the number, quality and mapping of the obtained reads is detailed in the Supplementary Table 2 for each omic approach respectively. Sequences used in this study are available at NCBI repository (BioProject ID: PRJNA795883).

**Figure 1.**
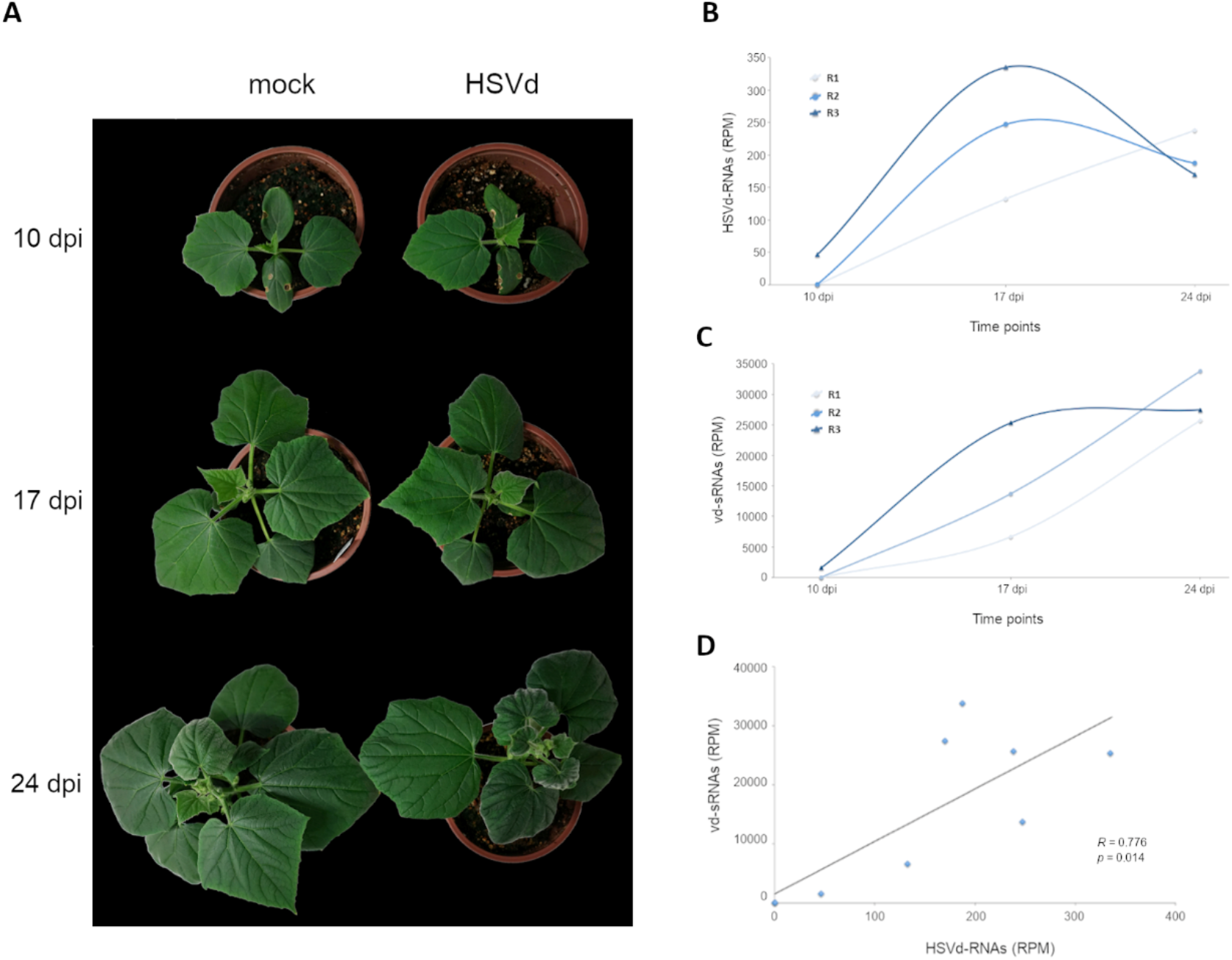
HSVd RNA is detected at 10 dpi in infected plants: **A**) Representative infected and mock inoculated cucumber plants at the three analyzed time points. Typical plant symptoms characterized by reduction in leaf size and incipient stunting are evident at 24 dpi. Graphic representation of the total transcripts (**B**) and sRNAs (**C**) derived from HSVd genome detected in apical leaves at 10, 17 and 24 dpi. **D**) Scatter plot showing the significant positive correlation (estimated by Pearson correlation coefficient) between the temporal accumulation of vd-transcript and viroid sRNAs (vd-sRNAs) in infected cucumber plants.

The infection rate of inoculated plants was estimated by considering the accumulation of viroid transcripts and viroid-derived sRNAs (vd-sRNAs) (Figure 1B and C). Viroid transcripts were detected in two (HSVd-R2 – 0.53 RPM and HSVd-R3 – 46.4 RPM) of the three analyzed samples at 10 dpi, while at 17 dpi an increased number of HSVd-related transcripts were recovered from all the analyzed samples (132.48, 247.13 and 335.04 RPM for replicates HSVd-R1, HSVd-R2 and HSVd-R3 respectively) (Supplementary Table 3). A lower accumulation (in comparison to 17 dpi) of viroid-transcripts was observed in HSVd-R2 (187.3 RPM) and HSVd-R3 (169.4 RPM) samples at 24 dpi. In contrast, the biological replicate HSVd-R1 (with no-detected viroid transcripts at 10 dpi), showed an increased accumulation (238.0 RPM) at 24 dpi (Supplementary Table 3). On the other hand, vd-sRNAs were only detected (1421.0 RPMs) in systemic tissues of the replicate HSVd-R3 at 10 dpi.

The accumulation of sRNAs homologous to HSVd in infected tissues at intermediate and late HSVd infection phases was more homogenous and vd-sRNAs were recovered from the three analyzed replicates at 17 dpi (6456.6 RPM, 13504.1 RPM and 25128.6 RPM) and 24 dpi (25539.9 RPM, 33627.7 RPM and 27284.7 RPM), revealing that vd-sRNAs increased through the analyzed period (Figure 1C).

In general, vd-sRNAs accumulation correlated (R = 0.776, *p* = 0.014) with the recovered HSVd transcripts (Figure 1D). HSVd transcripts and vd-sRNAs represented (in average) respectively, the 0.002% and 0.047% of the recovered reads at 10 dpi, the 0.024% and 1.5% at 17 dpi and the 0.02% and 2.9% at 24 dpi (Supplementary Table 3 and 4). Our results support that both, viroid transcripts and vd-sRNAs may be detected in systemic tissues in an irregular manner in early infection phases (10 dpi) and more consistently since the 17 dpi.

### Modifications in the host siRNA/miRNA population

Associations between sRNA expression profiles (considering control/infected plants and their biological replicates in the three time points) were evaluated using Principal Components Analysis (PCA). The percentage of total variance explained by the first three principal components (PC) was 45,91% and the treatments/time-points groups were significantly (*p* = 3.05’
s10 ^-12^) separated in the PC space (Supplementary Figure 2A). The endogenous cucumber sRNAs exhibited a distribution of read lengths enriched for 24 nt long (34.0%), followed by 23 nt (22%) and similar accumulations of 25 (14.3%), 21 (12.0%) and 22 (10.5%) nt long molecules. Reads of 20 nt represented the least abundant category (6.0%) (Figure 2B). This difference in accumulation of different sRNA lengths was statistically significant (2-ways non-parametric ANOVA, *p* < 2.12×10^−15^, 2.14×10^−13^ and 1.30×10^−15^ for 10, 17 and 24 dpi (respectively). In coincidence with diverse cucumber cultivars [54,55], this effect was predominantly due to the large enrichment in 24 nt long sRNAs in all pairwise comparisons. No significant differences were found between HSVd-infected and control conditions regarding the observed global distribution of sRNAs sizes (2-ways non-parametric ANOVA, *p* = 0.93; 0.85 and 0.55 for 10, 17 and 24 dpi, respectively).

**Figure 2.**
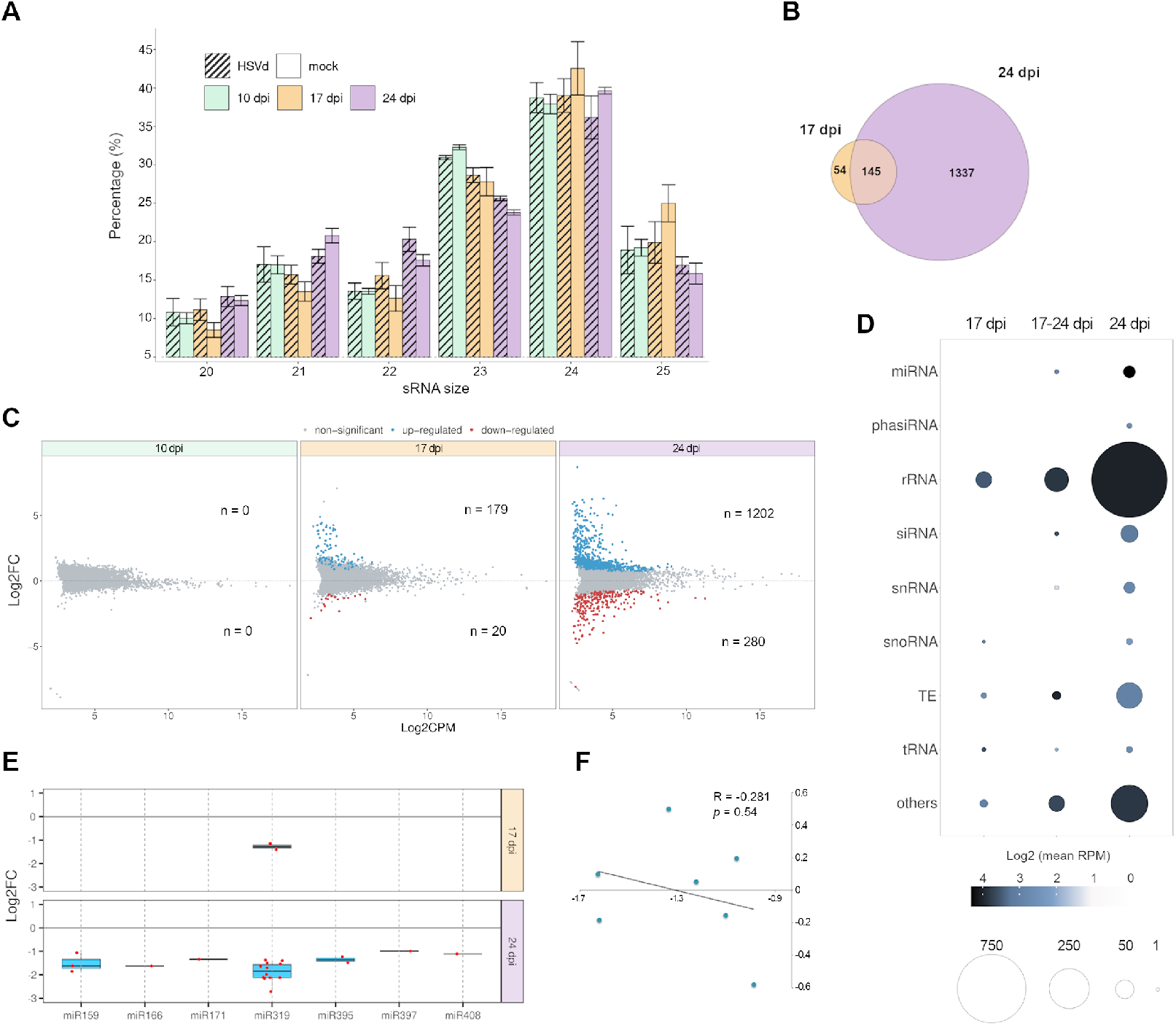
Analysis of the sRNA population recovered from the analyzed cucumber libraries. A) Diagram showing the means of the temporal relative accumulation and distribution of the total sRNAs reads ranging between 20 and 25 nt (Error bars indicate the SE between biological replicates). The three analyzed times are represented by colors (green: 10 dpi, orange: 17 dpi and magenta: 24 dpi). Smooth and striped bars represent data arising from mock-inoculated and HSVd-infected plants. **B**) Venn diagram representing the number of sRNAs responsive to HSVd infection at 17 and 24 dpi. **C**) Volcano plot representing the endogenous cucumber sRNAs with significant differential expression in infected plants at the three analyzed time points. Blue and red dots indicate up- and down-regulated sRNAs, respectively. The number of differential sRNAs in each time point is detailed. **D**) Categorization of HSVd-responsive sRNAs. The ball size represents the number of unique reads recovered. The color intensity indicates the sRNA accumulation estimated by the Log2 of the mean of the normalized reads. miRNA: micro RNA, phasiRNA: phased secondary small interfering RNA, rRNA: ribosomal RNA, siRNA: small interfering RNA, snRNA: small nuclear RNA, snoRNA: small nucleolar RNA, TE: transposable element, tRNA: transfer RNA. **E**) Box-plot analysis showing the general expression value observed for each miRNA-family member (dots) at 17 and 24 dpi. The differential expression of each miRNA family is represented by the median (internal box line) of the LFC values. **F**) Scatter plot showing the non-significant negative correlation (estimated by Pearson correlation coefficient) between the expression levels of the seven miRNAs responsive to HSVd infection and the accumulation of their targets in infected plants.

Only a reduced number (1536 unique reads) of endogenous sRNAs showed significant altered accumulation in HSVd-infected plants at 17 and 24 dpi (no differential sRNAs were detected at 10 dpi) (Figure 2C). 145 sRNAs were differentially expressed at both 17 and 24 dpi, while 54 and 1337 endogenous sRNAs, showed specific differential accumulation at 17 or 24 dpi, respectively. A total of 1246 (81.12%) and 290 (18.88%) sRNAs increased or decreased, respectively, their accumulation in response to infection. Under a temporal viewpoint, although no differentially expressed sRNAs were detected at 10 dpi, 199 reactive sRNAs (179 over-expressed and 20 down-regulated) were identified at 17 dpi (Figure 2D). The most evident alteration in the accumulation of endogenous sRNAs was observed at 24 dpi in which 1482 reads (1202 up-regulated and 280 down-regulated) were identified as differentially expressed in HSVd-infected plants. The 68.42% (1051 reads) of the differential sRNAs were categorized as derived from ribosomal RNA (rRNAs), while the 7.55% (116 reads) from transposable elements (TE), and the 3.06% corresponded to 47 reads identified as small interfering RNAs (siRNAs) previously described in cucumber (Figure 2E). A minor representation (1.5%) was observed for known microRNAs (miRNAs), sRNAs derived from snRNAs (1.3%), tRNAs (0.52%), snoRNAs (0.39%) and phased siRNAs (0.2%). The 17.06% of the sRNAs with differential expression were derived from unidentified or not representative functional categories.

We identified seven miRNA families (miR158, miR166, miR171, miR319, miR395, miR397 and miR408) with differential expression associated to HSVd-infection (Figure 2F). Except for miR319 (with 14 different sequences), reactive families were represented by one (miR166, miR171, miR397 and miR408), two (miR395) or three (miR159) unique sequences. All members in each miRNA family decreased in a coordinated manner in response to HSVd-infection (Supplementary Table 5). miR319 was the unique family that exhibit differentially expressed members at 17 and 24 dpi, the rest of the reactive miRNAs were only differential at 24 dpi. The differential expression values of the reactive miRNAs were low (in any case LFC > -1,6), suggesting that the regulatory pathways mediated by miRNAs are not particularly affected by viroid infection. The observation that such miRNAs negatively correlated in a non-significant manner (R = -0.281, *p* = 0.541) with their target transcripts reinforced this notion (Figure 2G).

### Characterization of of HSVd-derived sRNAs

A total of 1,523,822 reads (representing 7,428 unique sequences) fulfilling the conditions (detailed in material and methods) required to be unequivocally considered as HSVd-derived sRNAs (vd-sRNAs) were recovered from the three analyzed time points (Supplementary Table 3). Analysis of polarity distribution revealed a slight but significant (*p* = 2.2×10^−16^) difference according to the strand of origin of the vd-sRNAs (58.1% sense and 41.9% antisense) (Figure 3A). This polarity proportion was maintained constant in the three analyzed time points (Supplementary Figure 3). Considering each sRNA size-class individually, the sense-strand derived sRNAs were the predominant, except in 20 nt and 23 nt in length vd-sRNAs (Figure 3B). In contrast, a comparable accumulation rate for both plus and minus reads was observed when unique vd-sRNAs were considered (Figure 3C). Vd-sRNAs were mainly of 24 nt (68.9%, 57.1% and 49.0%,) and 21 nt (15.5%, 24.1% and 30.0%), for 10, 17 and 24 dpi, respectively. Except for 22 nt (10.3%, 11.8% and 12%), vd-sRNAs of 20, 23 and 25 nt accumulated under 5% of the total sRNAs homologous to HSVd in the three analyzed time points (Figure 3D). Regarding the predominant vd-sRNAs (24 nt and 21 nt) it was evident that while the proportion of 21 nt vd-sRNAs increases during viroid-infection, the accumulation of 24 nt reads recovered from infected plants decreases during the analyzed period. The observation that a comparable size-evolution pattern was not detected for endogenous sRNAs (Supplementary Figure 4) supports that this significant (2-ways non-parametric ANCOVA, *p* < 0.031 and 0.039, for 21 and 24 nt respectively) temporal bias in the accumulation of this type of vd-sRNAs is specific for the HSVd-RNA and not related to the cucumber sRNAs biogenesis pathway.

**Figure 3.**
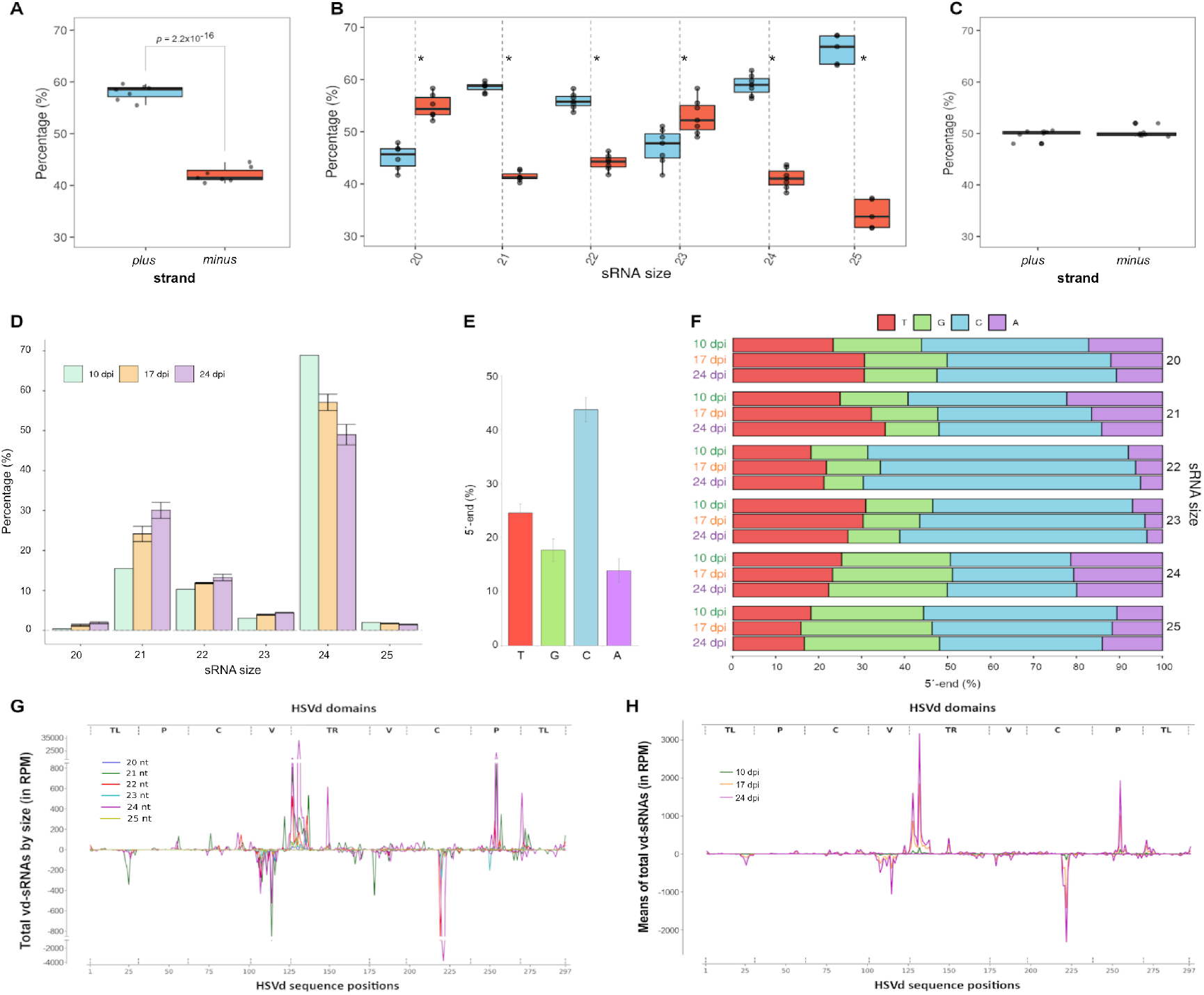
Characterization of vd-sRNAs. **A**) Box plot analysis of the relative accumulation of the total vd-sRNAs (20 to 25 nt) derived from genomic (plus) and antigenomic (minus) HSVd-RNA strand at the three analyzed time points. **B**) Distribution of polarity of the total vd-sRNAs discriminated by size, an asterisk indicates a p < 0.05. **C**) Comparative analysis of unique reads of vd-sRNAs. **D**) Diagram showing the means of the temporal relative accumulation and distribution of the vd-sRNAs reads ranging between 20 and 25 nts (the error bars indicate the SE). The three analyzed times are represented by colors (green: 10 dpi, orange: 17 dpi and magenta: 24 dpi). **E**) Graphic representation of the proportion of the 5’-ends in the total recovered vd-sRNAs. The means of the total reads are showed. **F**) Distribution of the 5’-ends discriminated by size and infection time. **G**) Genome view of the vd-sRNAs recovered from infected cucumber plants. The vd-sRNAs were plotted according to the position of their 5’
s-end onto the HSVd-RNA sequence in either sense (above the x-axis) or antisense (below the x-axis) configuration. The values on the y-axis represent the abundance of total vd-sRNAs (sizes indicated by colors) in the three analyzed time point. **H**) Representation of the temporal accumulation of total vd-sRNAs (means of total vd-sRNAs are represented). Vd-sRNAs recovered in each of the three analyzed times are represented by colors (green: 10 dpi, orange: 17 dpi and magenta: 24 dpi).

Categorized according to their 5’ terminal nucleotide, vd-sRNAs with a C-end were the most abundant (43,8%), followed by reads with T- (24,6%), G- (17,7%) and A- (13,9%) 5’
s ends (Figure 3E). Similar distribution of 5-ends was observed when total vd-sRNAs were analyzed considering size and temporal dynamics (Figure 3F).

When total vd-sRNAs recovered at the three time points were plotted onto HSVd-genome we observed that in general, sense and antisense vd-sRNAs spreading along the entire HSVd genome showed a heterogeneous distribution pattern with four evident hyper accumulation peaks in the position 130 and 254 and -78 and -184, in genomic and anti-genomic HSVd-RNA, respectively (Figure 3G). The hyper accumulation peaks predominantly corresponded to 24 nt sRNAs (Figure 3H). This global profile was maintained during the analyzed infection period (Supplementary Figure 5). In contrast to hyper accumulation peaks, we observed that a well-defined region of the viroid-genome comprised between the positions 11 to 38 (sense reads) and 31 to 58 (antisense reads) showed a null or much lower accumulation of vd-sRNAs in the three analyzed time points (Supplementary Figure 6).

### HSVd-infection modulates the expression and alternative splicing of host-transcripts

It is expected that the host transcriptome may change in response to the viroid infection. To test that assumption, RNA extracted from control and infected plants was subjected to RNA-seq. Associations between RNA expression profiles were evaluated using Principal Components Analysis (PCA). The percentage of total variance explained by the first three principal components (PC) was 62.52% and the treatments/time-points groups were significantly (*p* = 1.2×10^−10^) separated in the PC space (Supplementary Figure 2B). A high proportion (98.33%) of the recovered transcripts in both infected and control plants mapped to the cucumber genome (Supplementary Table 3), allowing the detection of differentially expressed transcripts (DETs) in all time points. The residual reads corresponded to unassigned transcripts (1.66%) and HSVd-derived sequences (0.01%). Considering only reads fully homologous to previously annotated cucumber RNAs, we identified 1125, 515 and 1390 transcripts with significant (FDR ≤ 0.05) differential accumulation in HSVd-infected plants, at 10, 17 and 24 dpi, respectively (Figure 3A and Supplementary Table 6). A total of 925 (at 10 dpi), 288 (at 17 dpi) and 1052 (at 24 dpi) host-transcripts presented time-specific differential accumulation (Figure 3B). Considering Log2FC ≥ 1 or ≤ -1 as cut-off value, the list of differential transcripts was reduced to 10 up-regulated at 10 dpi, 55 (20 up-regulated and 35 down-regulated) at 17 dpi and 271 (46 up-regulated and 225 down-regulated) at 24 dpi (size-increased dots in Figure 3A). The 10 up-regulated genes recovered at 10 dpi were time-specific, whereas 27 and 243 transcripts showed specific differential expression at 17 and 24 dpi, respectively (Figure 3C). Under a temporal viewing, it was obvious that the host transcripts with increased accumulation were predominantly (695 transcripts, 61.7%) recovered at 10 dpi, while down-regulation was predominant at 17 dpi (302 transcripts, 58.64%) and 24 dpi (975 transcripts, 70.14%). A similar trend was observed when expression values with a Log2FC ≥ 1 or ≤ -1 were considered. The amplitude (estimated by the variance of the differential expression values) and diversity (estimated by the number of DETs) of the transcriptional response to HSVd was clearly increased during the analyzed period of infection (Figure 3D).

According to the analysis of GO-terms, transcriptional response to HSVd-infection was characterized by a significant enrichment in genes associated to diverse biological categories mainly related to metabolic and cellular process, catalytic activity, ribosome metabolism, cellular/intracellular components, transport and membrane (Figure 4E and Supplementary Table 6). Under a global viewpoint, it was evident that the wide-range of enriched functional categories observed at 10 dpi (15 categories), was decreased during the infection (6 enriched-categories at 24 dpi). Searching for host genes potentially involved in a more persistent response to HSVd-infection we identified cucumber transcripts differentially expressed during the three infection phases. A total of 21 transcripts were identified as significantly responsive to infection at 10, 17 and 24 dpi (Figure 3F and Supplementary Table 7). Except for three specific transcripts up-regulated in at least one of the analyzed time points (Figure 3G, cluster 2), the common response of the other 18 differential transcripts to HSVd-infection was a stable (cluster 1) or temporally increased (cluster 3) down regulation. Genes included in this common response were functionally related to transport, membrane activity and response to biotic stimulus. This functional specialization was also supported by GO-terms enrichment when were considered only transcripts with Log2FC ≥ 1 or ≤ -1 with common differential expression at late (17 and 24 dpi) infection phases (Figure 4H). Differential transcripts without previous annotation (and consequently not considered in our study) were analyzed as is described in Methods section (Supplementary Figure 7). The obtained results revealed that such transcripts (mainly identified as non-coding RNAs) exhibited minimal alterations in response to HSVd infection (Supplementary Table 8).

**Figure 4.**
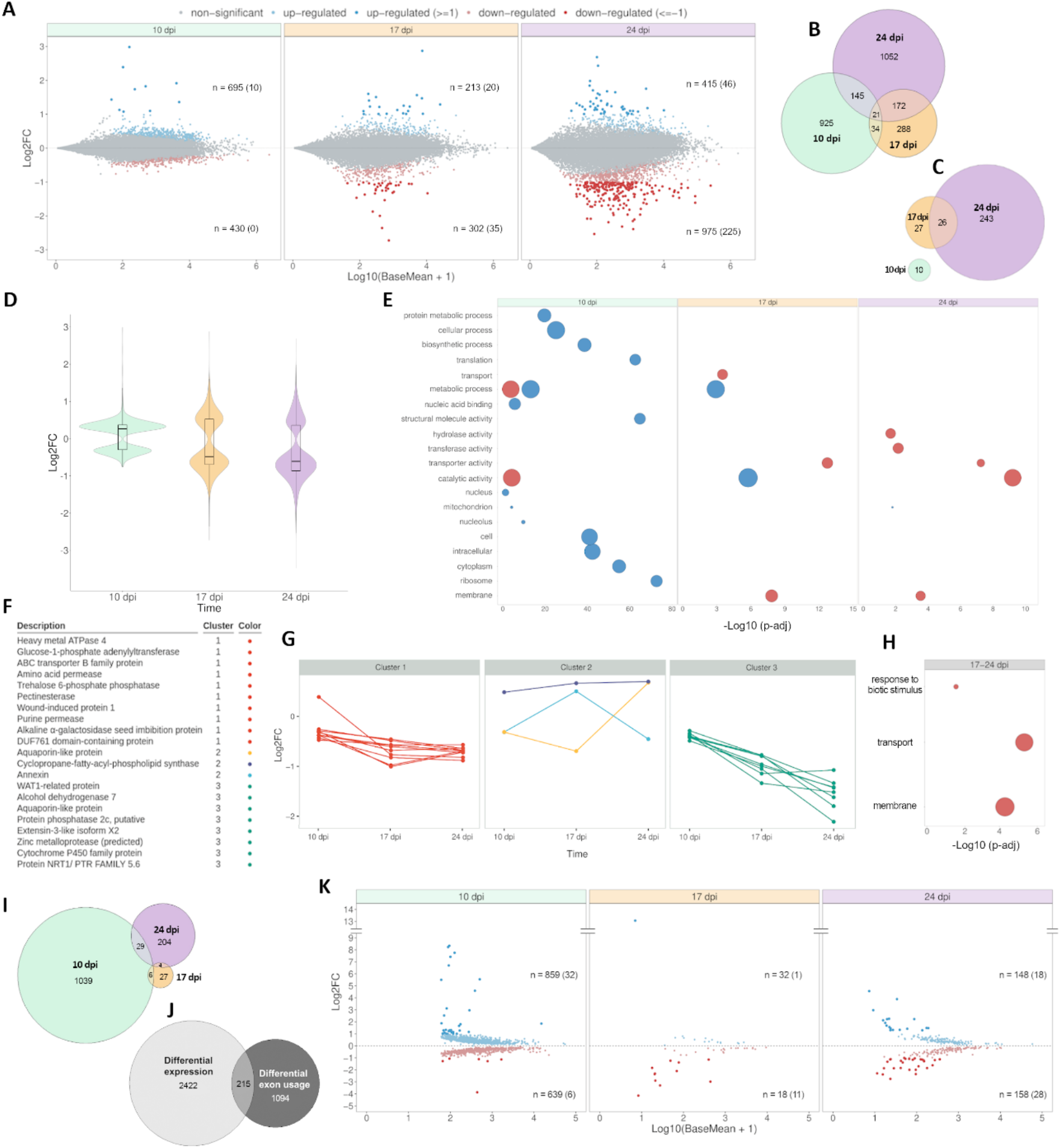
Host-transcriptional alterations associated to HSVd-infection. **A**) Volcano plot representing the differentially expressed transcripts (DET) in infected cucumber plants at 10, 17 and 24 dpi. Blue and red dots indicate up- and down-regulated transcripts, respectively. DET with LFC ≥ 1 or ≤ -1 are represented by bold dots and the number is indicated in brackets. Venn diagram showing all the DET (**B**) and those with LFC ≥ 1 or ≤ -1 (**C**) identified at 10, 17 and 24 dpi **D**) Violin plot representing the temporal profile of the transcriptional alteration associated to HSVd-infection. Internal box line indicates the median of the LFC values. **E**) Gene ontology analysis (plant GOSlim) for DETs identified in HSVd-infected plants at 10, 17 and 24 dpi. Circle size is proportional to the fraction of genes relative to the total number of genes with the GO term. Color indicates whether the genes are upregulated (blue) or downregulated (red). The –Log(10) of the adjusted P values is represented in the X-axis. **F**) Detail of the 21 transcripts differentially expressed at the three analyzed time-points. **G**) Clustering analysis of time-course expression profiling of transcripts with differential expression at 10, 17 and 24 dpi. **H**) Gene ontology analysis (plant GOSlim) for transcripts differentially expressed at both 17 and 24 dpi (LFC ≥ 1 or ≤ -1). **I**) Venn diagram showing the number of differential exon usage (DEU) events identified at 10, 17 and 24 dpi and (**J**) transcripts showing common and specific regulation by differential expression or exon usage. **K**) Volcano plot representing the events of DEU in infected plants at 10, 17 and 24 dpi. Blue and red dots indicate up- and down-regulated exons, respectively. Events of DEU with LFC ≥ 1 or ≤ -1 are represented by bold dots and the number is indicated in brackets.

Alternative splicing (AS) has been proposed as a regulatory mechanism crucial for the modulation of the plant development and response to virus infection [56]. Sequenced transcripts were analyzed with DEXSeq tool to infer the AS landscapes of cucumber plants and determine their differential patterns during HSVd infection. We identified 1589 differential exons derived from 1309 intron-containing multiexonic genes that were alternatively spliced in infected plants (Figure 3I). Among such transcripts, only 215 (5.76 %) also showed differential expression (Figure 3J). Under a global viewpoint, significant differential exon usage (DEU) was predominantly observed in infected plants at 10 dpi (1498 exons with DEU), less frequent at 24 dpi (306) and a residual phenomenon at 17 dpi (with 50 alternative splicing events identified) (Figure 3K and Supplementary Table 9). Differential exon usage in response to HSVd-infection was comparable at early (38 exons) and late infection phases (46 exons) when we considered only Log2FC values ≥ 1 or ≤ -1 (size increased dots in Figure 3K). Regarding the temporal trend in the differential exons usage, our results support that HSVd infection is associated to a predominant retention of exons at 10 dpi showing 32 over-represented exons with Log2FC values ≥ 1 (Figure 4K). In contrast, the underrepresented exons (with Log2FC ≥ -1) were prevalent at 17 (11 exons) and 24 dpi (28 exons). Transcripts exhibiting frequent DEU, at both early (10 dpi) and late (24 dpi) infection phases were predominantly involved in primary metabolism-related process (Supplementary Figure 8). Additionally, transcripts associated to membrane and transport, were enriched at 10 dpi.

### Epigenetic alterations associated to HSVd-infection

To analyze if HSVd-infection modifies the epigenetic landscape of cucumber plants, whole-genome bisulfite sequencing (WGBS) libraries were constructed. Total sequence-coverage obtained from both mock and HSVd-inoculated plants was comparable attesting for the reproducibility of our assays (Supplementary Figure 9). Global analysis (considering the total percentage of cytosine methylation in CG, CHG and CHH sequence context) revealed a non-significant hypomethylation in infected plants at 10 dpi (70.39% and 67.21%, for control and infected respectively), followed by a slight hypermethylation at 17 dpi (75.04% and 76.93%) that was significant (70.6% and 721% for control and infected plants, respectively) at 24 dpi (Figure 5A, upper panel). These differences were maintained when we considered total methylation level for each sequence context specifically (Figure 5A, lower panel). A similar scenario was observed when the total methylation levels were analyzed, considering each cucumber chromosome individually (Figure 5B). To obtain a more precise picture of the specific methylation changes, we used DMRcaller to identify differentially methylated regions (DMRs) for the three CG, CHG, and CHH sequence contexts. Only significant DMRs with differences in methylation ≥15% were considered. The hypomethylated CG DMRs were the most abundant during the analyzed infection period. In contrast hypermethylated DMRs were most abundant considering CHG and CHH sequence context (Figure 5C).

**Figure 5.**
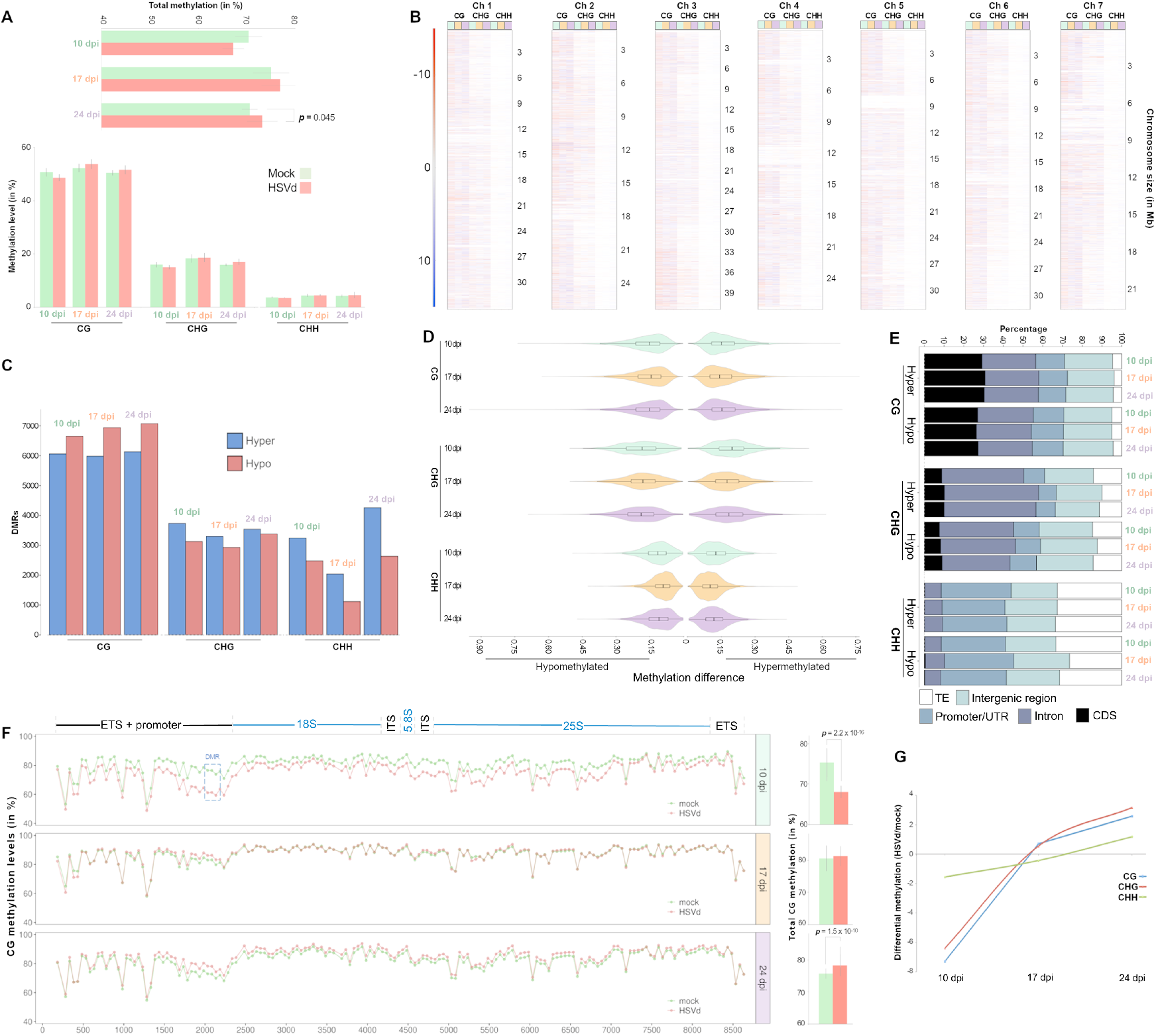
HSVd induces alterations in the cucumber epigenetic landscape. **A**) Graphic representation of the total cytosine methylation mean values of HSVd-infected and control plants at 10, 17 and 24 dpi (upper panel) and discriminated by sequence context (lower panel). Error bars indicate the standard error between replicates. **B**) Chromosome view of the temporal dynamics of the differential methylation profiles in HSVd-infected plants at 10 (green), 17 (orange) and 24 dpi (purple). Red and blue lines represent hypomethylated and hypermethylated regions, respectively. **C**) Number of significant hyper- or hypo-differentially methylated regions (DMRs) in the three sequence contexts (CG, CHG and CHH) identified in infected plants at 10, 17 and 24 dpi. **D**) Violin plot representing the temporal profile of the epigenetic alterations (estimated by significant DMRs) associated to HSVd-infection. Internal box line indicates the median of the percentage of differential methylation values. **E**) Temporal description of the proportion of the cucumber genome regions containing significant DMRs identified in infected plants. **F**) Global view of the methylation profiles in CG context in ribosomal RNA transcriptional unit at 10, 17 and 24 dpi in HSVd-infected (red) and control (green) plants. In the left part each dot represents the mean methylation values of 50 nucleotides while in the right the total mean methylation values of the whole ribosomal unit are represented. A significantly hypomethylated DMR identified in infected plants at 10 dpi and matching to promoter region is highlighted. The ribosomal genes (18S, 5.8S and 25S) are indicated in blue while the internal transcribed spacers (ITS) and external transcribed spacers (ETS) are in black **G**) View of the temporal dynamics of the differential methylation in the three sequence contexts (CG, CHG and CHH) observed in ribosomal RNA transcriptional unit during the HSVd infection.

This global trend was maintained constant during the three analyzed time points. Considering the intensity (assumed as the median of the DMRs values) of the methylation alterations in each time and context, it was evident that the higher differential values for both hypo (18.4%, 18.0% and 18.8% for 10, 17 and 24 dpi, respectively) and hypermethylation (19.8%, 17.5% and 18.3%10, 17 and 24 dpi, respectively) were observed for the GHG context, while a lower number of alterations was observed for the CHH DMRs (11.3%, 9.0% and 10.8%) for loss of methylation at 10, 17 and 24 dpi, respectively and 12.5%, 9.9% and 11.6% for gaining in identical time points (Figure 5D). For the DMRs of the CG context, the medians of the differential alterations were near to the 15% during the entire analyzed period. Considering the functional categories of the genomic position of the DMRs, approximately 60% of the observed CG DMRs in the three analyzed time points were mapped in regions predicted as coding sequences (CDS) and introns, while intronic regions constituted the predominant DMRs at the CHG context (Figure 5E). As expected, during the analyzed period, CHH DMRs were enriched in TEs and strongly diminished in CDS.

It has been previously demonstrated that HSVd infection promotes alterations in the epigenetic landscape of regulatory regions of the ribosomal DNA (rDNA) [47] to reconfigure a functional scenario that encourages its replication in cucumber plants [48]. To extend this study, we analyzed if rDNA was specifically identified as a significant DMR. A region of 150 nt comprised between positions 1984 – 2133 of the ribosomal genes and overlapping with their promoter region was identified as hypomethylated DMR in CG and CHG contexts at 10 dpi (boxed region in Figure 5F and Supplementary Figure 9). Moreover, considering the total methylation levels at CG context, it was evident that HSVd induces a significant (*p* = 2.2×10^−16^) hypomethylation of the rDNA (79.57 and 775 of total methylation for mock and HSVd-infected plants, respectively) at 10 dpi. Although the downregulation of the methylation levels was more evident in the promoter region (identified as a significant DMR), the hypomethylated status was extended to the entire ribosomal gene (Figure 5F – upper panel).

Epigenetic alteration of the rDNA associated to HSVd infection was revealed as a dynamic phenomenon, being comparable to mock at 17 dpi (Figure 5F – central panel) and significantly (1.5×10^−10^) hypermethylated at 24 dpi with total-CG methylation levels of 80.94% and 837% for mock and HSVd-infected plants, respectively (Figure 5F – lower panel). This dynamic alteration of the host epigenetic landscape was also observed when CHG and CHH sequence contexts were considered (Figure 5G and Supplementary Figure 10). Under a temporal viewpoint, the significant hypomethylation of ribosomal genes was coincident with the initial phase in which HSVd reaches a systemic infection, while the transition to an hypermethylation status in host rDNA was in parallel with the stabilization (or slight decrease) of the HSVd transcripts in infected plants (Supplementary Figure 11).

### Interplay between methylation and transcriptional regulation in infected plants

To analyze if the changes in the methylation levels observed in HSVd-infected plants could be associated to transcriptional alterations we integrated the data obtained by RNA-seq and WGBS assays. 113 protein-coding transcripts with strong (Log2FC ≥ 1 or ≤ - 1) differential expression at 24 dpi containing at least one DMR in any of these regions (promoter/UTR, CDS and intron), were selected for this analysis (Supplementary Table 10). We observed that 68 of these 113 genes —representing a significant (*p* = 0.038 in Exact binomial test) proportion (60.2%) of the analyzed transcripts— were characterized by containing at least one antagonistic DMR (hypomethylated for over accumulated transcripts or hypermethylated for downregulated transcripts) (Supplementary Table 11). Interestingly, a significant negative correlation was observed when we compared their expression levels (estimated by Log2FC) with their global methylation status (determined by DMR analysis in the three sequence contexts) in promoter/UTR and CDS regions (*p* = 0.003 and 0.014, respectively) (Figure 6A). No significant correlation was observed for DMRs matching to introns (*p* = 0.148). A more detailed analysis (focused on each context separately) evidenced that the negative correlation was mainly associated to DMRs in the CG context (Figure 6B and Supplementary Figure 12). Next, we compared the expression and methylation levels of these 68 genes at 10, 17 and 24 dpi to obtain a global overview of the temporal dynamics of the interplay between cytosine methylation and transcriptional response during HSVd-infection. Our results indicate that while the intensity and the density of the global methylation of the analyzed genes (estimated by the mean values and the total number of the DMRs) were increased during the analyzed period (Figure 6C, upper panel - left), their transcriptional activity (estimated by the mean of the Log2FC values and the number of DET) was consistently down-regulated (Figure 6C, lower panel - left). In a similar way, the transcripts upregulated during HSVd-infection were associated to a temporally increased hypomethylation status of their respective genes (Figure 6C, right - lower and upper panel respectively). According to their biological function, the cucumber genes showing methylation levels antagonistic to transcripts accumulation mainly encoded proteins related to membrane (associated to stress response or transport), oxidation-reduction and metabolic processes (Supplementary Table 11). Additionally, some genes encoding for nucleus components, transcription factors and protein involved in protein-protein interactions, were also identified.

**Figure 6.**
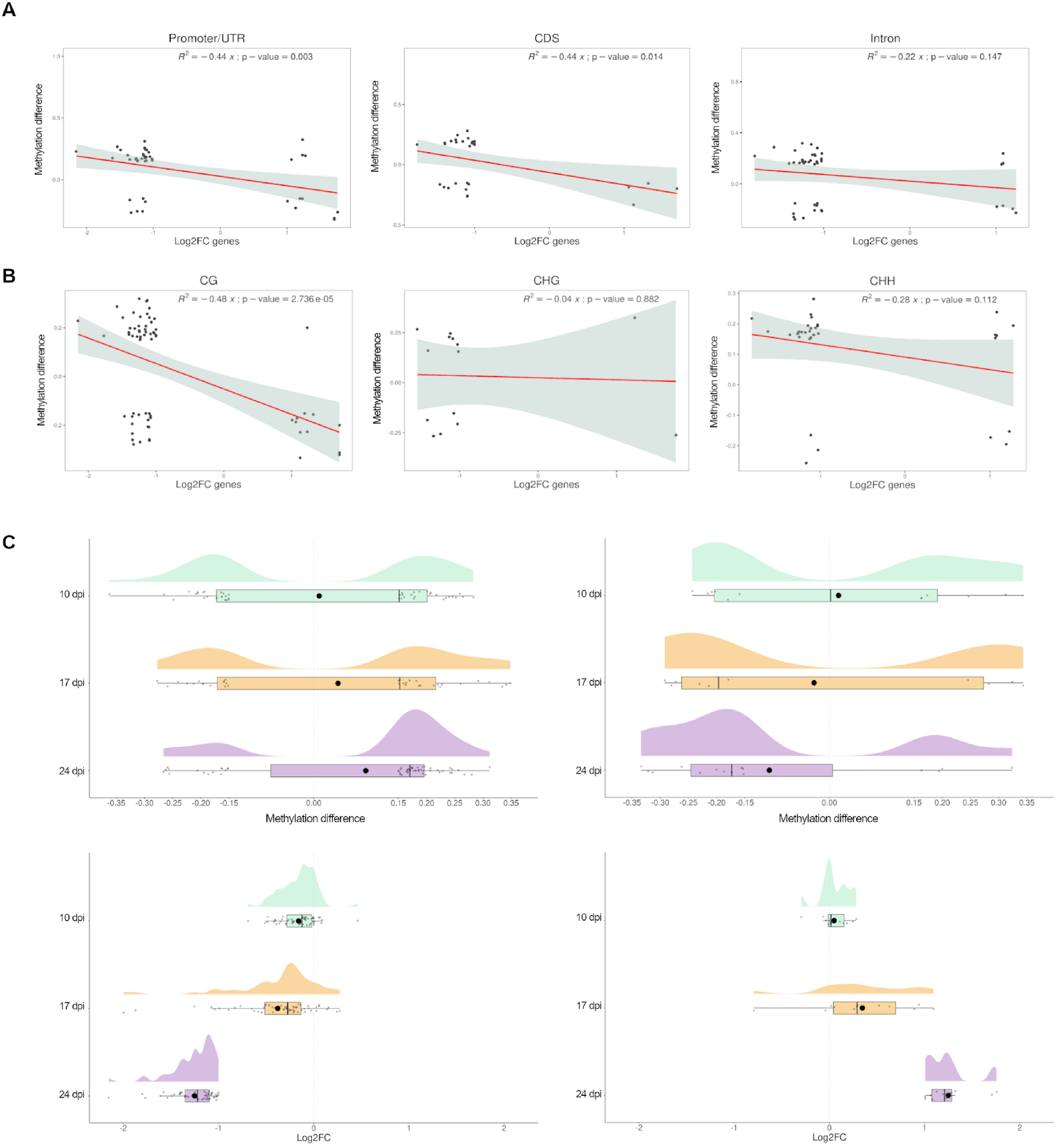
Association between transcriptional alterations and epigenetic changes induced by HSVd-infection. Scatter plots showing the negative correlation (significance estimated by Pearson correlation coefficient) between the expression levels of differential genes containing at least one antagonistic DMR and their global methylation status. **A**) Considering the three selected gene regions (Promoter/UTR, CDS and intron) and **B**) considering methylation sequence context. **C**) Raincloud plots showing the temporal correspondence between the sense and the intensity of the epigenetic changes (upper panels) and the transcriptional activity (lower panels) in response to HSVd-infection. The temporal increase in the intensity of the hypermethylated (upper left) and hypomethylated (upper right) DMRs represented by the mean of the differential methylation (black dots) is associated with an evident decrease (lower right) or increase (lower left), respectively (in intensity) of the transcriptional activity estimated by the LFC mean values (black dots).

Regarding the regulation of non-coding transcripts by epigenetic changes, we analyzed the transcriptional activity of the cucumber ribosomal genes, which has been earlier described to be modulated by HSVd infection [47]. Our data showed that the significant hypomethylation of rDNA observed at early infection phases was associated to an increased transcriptional activity of rRNA genes determined by the increased accumulation of primary precursors of the ribosomal transcripts (pre-rRNAs) in infected plants (Supplementary Figure 13, left panel). During rRNA transcription, RNA polymerase I transcribes length-units pre-rRNAs that are extensively processed into 18S, 5.8S and 25S units by the sequential deletion of external and internal transcribe spacers (ETS and ITS, respectively) [57]. Consequently, the differential accumulation of pre-rRNAs was estimated by considering the transcripts matching to ITS1 and ITS2 regions. Finally, the over accumulation in infected plants of rRNA-derived sRNAs (rb-sRNAs), an indirect indicator of the transcriptional activation of normally inoperative rRNA transcriptional units [58], provided additional support to the link between rDNA hypomethylation and transcriptional deregulation (Supplementary Figure 13, right panel).

## DISCUSSION

In coincidence with other cellular parasites, viroids are compelled to subvert all sorts of cellular factors and reprogram host gene expression in order to modulate (into their own benefit) complex plant regulatory networks. Assuming this functional scenario, it is expected that the type of affected host-regulatory pathways as well as the intensity of the induced alterations may vary considerably during the infection process. However, our knowledge about the functional diversity and temporal dynamics of the global plant response to viroid infection is limited [5,7].

Here, we have addressed this question by means of a temporal analysis of the global response to HSVd infection focused on three host regulatory levels small RNA interference, modulation of the transcriptional activity and epigenetic modifications. This integrative analysis allowed us to determine that during the analyzed infection period cucumber plants follow different strategies to modulate their cellular homeostasis in response to the functional alterations induced by HSVd replication and accumulation.

The early detection (at 10 dpi) of viroid-derived transcripts and sRNAs in non-inoculated apical tissues evidenced the capacity of HSVd to develop a relatively quick systemic infection in cucumber plants. However, the observation that viroid transcripts are recovered from only two out the three biological replicates and in a heterogeneous level (27 and 2487 and 2.94 and 1421, normalized reads, for genomic and vd-sRNAs respectively) suggests that the initial infection phase is highly variable and may be dependent on yet unknown viroid-host interactions or that minimal variations on the plant phenological stage could condition these early events.

Analysis of the small RNA data evidenced that, although sRNAs derived from *plus* polarity transcripts were predominantly recovered from infected plants at the three analyzed time points, both *plus* and *minus* HSVd replication intermediates are potential substrates for host RNA silencing machinery at different infection phases. In fact, considering the number of unique sequences no differences were observed between the two polarities. Our results are in consonance with previous studies in which the recovery of small RNAs of both polarities supports the predominant involvement of viroid replication intermediates as source of vd-sRNAs in plants infected by both nuclear [16,17,59–61] and chloroplastic [15,62] viroids. Considering the size distribution, 24 nt vd-sRNAs were predominant at the three analyzed time points, followed by 21 nt sequences. However, it was evident that this size-distribution profile was modified during HSVd infection, being characterized by a temporal increasing of the global proportion of 21 nt that negatively correlates with the accumulation of 24 nt vd-sRNAs. These results support that the processing/accumulation of 24 nt vd-sRNAs is predominantly associated to initial infection steps, while the accumulation of the 21 nt class constitutes an event characteristic of well-established infectious processes. The observation that a similar scenario is not observed when cucumber sRNAs were considered evidenced the independence of this phenomenon with the natural evolution of the endogenous sRNAs biogenesis. This temporal trend may explain the inconsistency between our data and previous results supporting that 21 nt in length is the most abundant class of vd-sRNA in HSVd-infected cucumber plants at late infection phases [60,63]. When the distribution of vd-sRNAs along the HSVd-genome was analyzed, we observed that reads matching onto four specific regions represent approximately the 40% of the recovered reads in the three analyzed time points. In contrast, two HSVd-RNA regions exhibited null or extremely lower accumulation of vd-sRNAs. The existence of these hyper- and hypo-accumulating regions (detected in both HSVd strands) may be explained assuming the existence of certain viroid-RNA regions exhibiting differential accessibility to host RNA silencing machinery and/or differential stability rates for these vd-sRNAs. It is interesting to note that one of these hypo-accumulating regions, the terminal left (TL), contains the terminal conserved hairpin (TCH), a highly conserved region in the HSVd sequence [64] and that besides *Hostuviroid* is also present in *Cocaviroid* and *Coleviroid* genera. In the *Pospiviroid* genus, this region harbors the DNA-dependent RNA polymerase II binding region and the transcription initiation site [65].

Having established the global landscape of pathogen-derived sRNAs, next we analyzed the influence of the HSVd infection in the endogenous sRNA population. A first view evidenced that the general profile of cucumber sRNAs is not affected in a significant manner in any of the analyzed time points. Considering sequence accumulation specifically, we observed that the biogenesis of endogenous sRNAs remains relatively stable in response to viroid accumulation, with only 1482 unique sRNAs with significant altered expression at 24 dpi. The predominant trend of this response was the upregulation of the host-derived sRNAs. Differentially recovered sRNAs were not identified at 10 dpi, while a considerable proportion of the sRNAs with significant differential expression observed at 17 and 24 dpi were derived from ribosomal transcripts and associated to hypomethylation of rDNA induced by HSVd-infection (discussed in depth below). A similar scenario (characterized by a lower reactivity level) was observed when alteration in miRNA population was analyzed. Seven miRNA families were significantly down-regulated in infected plants, and except for miR319 (responsive also at 17 dpi) the remaining miRNAs were differentially expressed only at 24 dpi. The association between miR319 downregulation and infection was also evident considering the number of family members (12 at 24 dpi) that showed comparable differential expression. However, the changes observed in miR319 level were not associated to antagonist alterations in the accumulation of their regulatory targets. It is well established that TEOSINTE BRANCHED1/CYCLOIDEA/PROLIFERATIN CELL FACTOR (TCP) mRNA is a highly conserved target of miR319 in diverse plant species [66] and accumulating evidence has revealed that miR319-regulated TCP (MRTCP) genes participate extensively in plant development by regulating hormone metabolism mediated signaling [67]. Interestingly, it has been previously reported that tomato plants infected with PSTVd (another member in the *Pospiviroidae* family) exhibit a complex array of changes affecting hormone signaling [30]. However, the lack of transcriptional alterations in MRTCP genes argues against the possibility that such physiological effects can be associated to HSVd infection in cucumber. No significant correlation was observed when the targets of the other six differentially expressed miRNAs where analyzed evidencing that, in coincidence with the described in other plant-viroid models [27,31,68,69], the global effects of HSVd infection in cucumber miRNA metabolisms is weak, and generally not related to an evident regulation of their targets.

Since the first viroid was described, it has been proposed that this type of naked exogenous RNAs may act as an abnormal regulator affecting host transcriptional activity [70]. Furthermore, the alteration of the accumulation of plant transcripts, at specific infection phases, has been previously reported in diverse viroid-host interactions [30,38,40,69,71]. Our temporal analysis of the evolution of the cucumber transcriptional landscape in response to HSVd infection indicates that although significant transcriptional alterations were observed in the three analyzed time points, the intensity of this response (considering both, the number of differential transcripts and the alterations in the accumulation levels) paralleled the development of the infection, being slight at 10 dpi and more evident at 24 dpi. Another particularity associated to the infection time was the trend of the transcriptional response, while the upregulation of host transcripts was the predominant response at 10 dpi, the downregulated transcripts were predominant at 17 and 24 dpi. According to the functionality, it was evident that HSVd-infection is associated to an extensive reprogramming of the host transcriptional activity, affecting cucumber genes involved in diverse cellular functions. This observation is in agreement with previous data obtained from plants infected with two (severe and mild) HSVd-variants, showing a complex array of changes in the host transcriptome associated to viroid infection [42]. However, a more defined functional response to HSVd-infection was evident when we considered only transcripts differentially expressed at 17 and 24 dpi. Cucumber genes included in this common response were predominantly downregulated and mainly involved in membrane metabolism, transport and response to biotic stress.

Besides transcript accumulation, it is well established that alternative splicing is a regulated process that increases the transcriptome diversity, providing an alternative way to favor the plant adaptation to environmental changes [72,73]. However, our knowledge about the alternative processing of host transcripts in response to viroid infection is very limited and only data related to PSTVd-infected tomato plants at late infection has been reported [69]. Our results evidenced that the differential exon usage was the predominant transcriptional response during the initial phase of the HSVd-infection (10 dpi). In contrast to the observed for transcripts expression, the global trend (up or downregulation) in exon accumulation was comparable at the three analyzed time points. Only a small proportion (5,76%) of the transcripts having AS, also exhibited significant differential accumulation, suggesting that the control of the transcriptional activity and the alternative processing of pre-mRNAs might be complementary mechanisms activated in cucumber plants in response to HSVd infection.

Regarding to the functionality of the transcripts showing significant AS in response to HSVd-infection, we observed that although the initial response included cucumber genes involved in many cellular processes, transcripts showing a more constant differential exons usage frequency during infection were predominantly related to primary metabolism. The observation that the AS of transcripts with similar biological functions was also reported in PSTVd-infected tomato plants [69], suggest that the regulation of the plant metabolism mediated by differential exon usage might be a host response mechanism also extended to other nuclear-replicating viroids.

The differential methylation of cytosines is another host regulatory layer susceptible to be affected in response to viroid infection [74]. Although, this epigenetic mechanism was first described studying viroid infection in PSTVd-expressing transgenic tobacco plants [75], the global effects of viroid-infection into host DNA methylation, remain yet unexplored [7]. The WGBS analysis evidenced that HSVd infection is associated with significant alterations in the host epigenetic landscape. Under a global viewpoint, the alteration in cytosine methylation was a dynamic phenomenon associated to infection phases (slight global hypomethylation at early infection and significant hypermethylation at 24 dpi) and sequence context (predominant hypomethylated DMRs at CG context and hypermethylated CHH). Studying in detail the regulatory effects of this altered epigenetic landscape; we observed that the changes in methylation levels (at CG sequence context) of putative promoter regions and CDS correlate with the transcriptional alterations observed in infected plants. Thus, suggesting that these HSVd-associated epigenetic variations may be responsible of the differential expression of certain genes in infected cucumber plants. Previously, it was reported that in cucumber [47,49] and *N. benthamiana* [76] plants, HSVd accumulation is associated to significant hypomethylation of the promoters region in rRNA genes. The results obtained here, besides reinforcing the previous results, demonstrated that the alterations in rDNA methylation are also extended to the totality of the rRNA transcriptional unity which is in coincidence with the global trend in the whole genome.

The integrative analysis of our data supports that in response to HSVd significant alterations are triggered in cucumber plants at different regulatory levels that are closely related to the temporal dynamics of the infection process (Figure 7). Differential exon usage, modulated by alternative splicing, emerges as the predominant response at the initial infection stage. It is well established that the alternative processing of the pre-mRNAs increases the diversity of the plant transcriptome and proteome [72]. Consequently, this rearrangement of the host-transcriptome (without new transcriptional activity) might, in similar way to the previously described for other stress conditions [73], allow the immediate fine tuning of the host gene expression in response to HSVd-infection. At subsequent infection phases, the modulation of the plant-response to the viroid will be mainly promoted by alterations in the transcriptional activity that was evidently increased at 17 dpi and being the mostly affected regulatory-layer (considering both, intensity and diversity) at 24 dpi. Interestingly, altered transcription shows a temporal coincidence with changes in the methylation profiles of promoter regions in certain cucumber genes. This functional connection permits to speculate about the possibility that the non-immediate transcriptional response to HSVd might be modulated by the host epigenetic changes associated to infection. Finally, it was obvious that the alteration in sRNA and miRNA metabolism played a minor functional role in the recovering of the cell homeostasis in response to HSVd infection.

**Figure 7.**
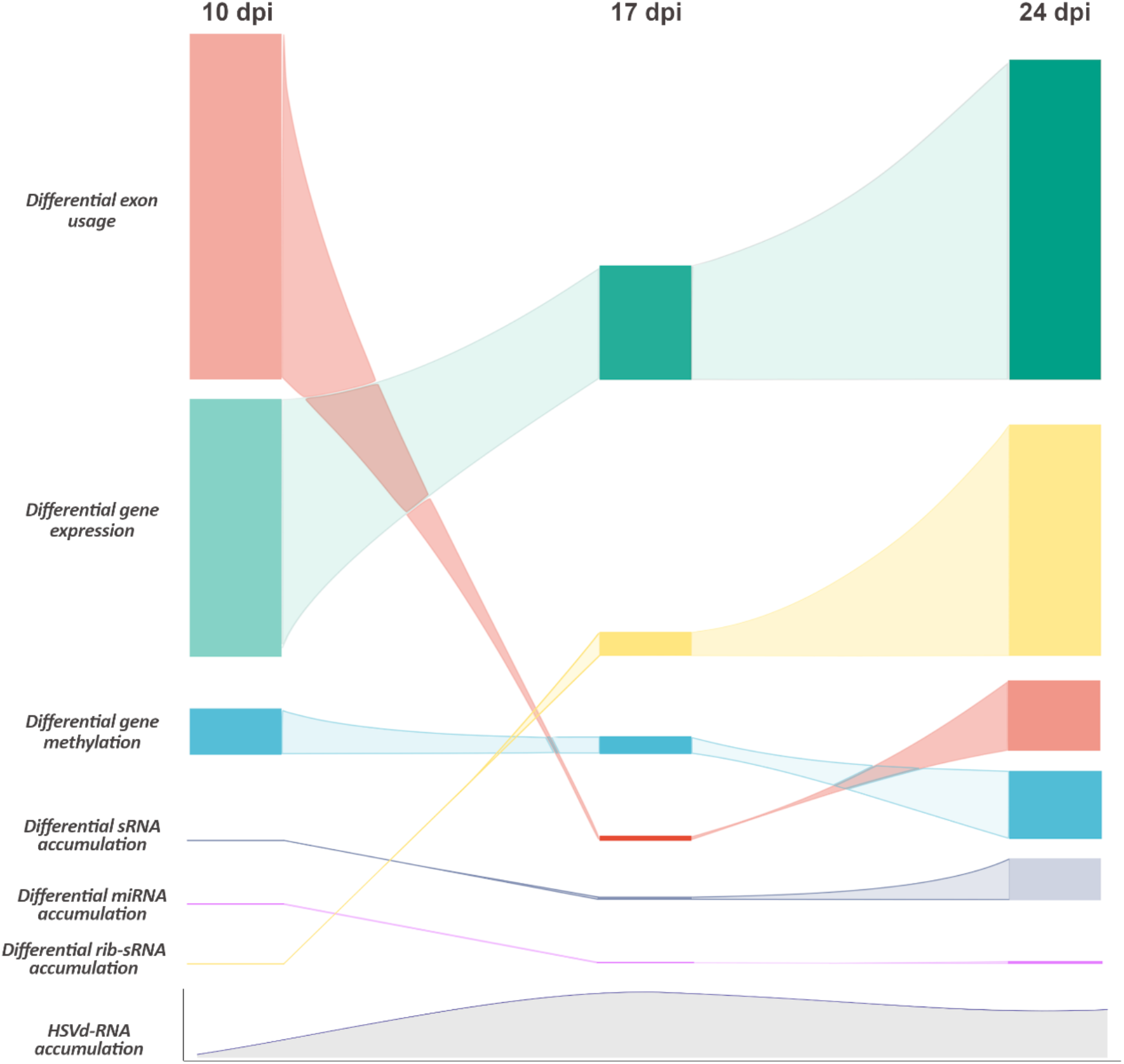
Graphic representation showing the temporal dynamics of the global host-regulatory response to HSVd-infection in cucumber plants. The predominant regulatory layers affected in each analyzed infection time are hierarchical represented by the density (box length) estimated by the number of alterations and intensity (box color strength) estimated by the absolute differential value of the response. To represent the density of the alterations in methylation we considered the number of genes with significant DMRs in CG context. In the sRNA group are included all the identified classes except rib-sRNAs and miRNAs. The global position in each analyzed time-point indicates the predominance of the regulatory layer in the global response to infection. Below it is also represented the relative accumulation (estimated by the amount of HSVd transcripts) in infected plants.

An interesting emerging question is how HSVd may trigger the regulatory alterations observed in infected plants. It is generally proposed that disease development must be the result of the direct interaction between viroid-RNA and certain host factors [5]. For example, it has been recently shown that the direct interaction between two symptomatic viroids (PSTVd and CEVd) and the host translational machinery induce ribosomal stress in tomato plants [77]. Assuming this possibility, it may be hypothesized that resembling the described for the arabidopsis long ncRNA ASCO [78], HSVd (an exogenous lncRNA) might bind to specific splicing factors affecting pre-mRNA processing. Regarding transcriptional alterations linked to host epigenetic changes, it has been proposed that the direct interaction between viroid-genome and HISTONE DEACETYLASE-6 (HDA6) is responsible of the methylation changes observed in the rDNA of HSVd-infected cucumber plants [48]. Consequently, the possibility that the HSVd might modulate the host epigenetic landscape by sequestering key components of the RNA-directed DNA Methylation (RdDM) pathway should not be yet excluded. Alternatively, the specific silencing of certain host genes mediated by vd-sRNAs can also contribute to the global redesign of the host regulatory structure. Further studies focused on a deep analysis of the potential interactions between viroid-RNA and host components will be needed for understanding the ways employed by HSVd to redesign the cucumber regulatory scenario.

Altogether, the data showed here constitute a comprehensive multiomic approach aimed to provide an overview of the temporal dynamics of the host response to HSVd infection. Recognizing that this is only the initial step in a long road, we expect that this innovative viewing will contribute to elucidate the mechanistic and the molecular basis of the host changes triggered by viroid infection.

## MATERIAL AND METHODS

### Viroid infection and sample collection

Cucumber plants cv. Marketer at cotyledon stage were inoculated with *Hop stunt viroid* (Y09352.1) as previously described [79]. Both cotyledons were agro-infiltrated with a culture of *Agrobacterium tumefaciens* strain C58 with 1 of optical density, harbouring pMD201t-HSVd or empty vector, diluted in infiltration buffer (MES 0.1 M, MgCl2 0.1 M). Plants were kept in a photoperiod of 16 h under visible light and 30 °C (light)/25 °C (darkness). The second leaf after the apex was collected at 10-, 17- and 24-days post inoculation. Each bio-replicate consisted of the leaves from three plants and for each time-point three bio-replicates were sampled of mock and HSVd infected plants.

### DNA and RNA extraction and library preparation

Total RNA was extracted using TRIzol reagent (Life Technologies) following the manufacturer instructions. Genomic DNA was extracted using a CTAB-based extraction method as previously described [80]. All libraries were constructed by Novogene Europe (Cambridge, U.K.) according to their standard procedures, including the previous steps of mRNA purification with oligo(dT) beads sRNA purification by size selection.

### Small RNA sequencing and analysis

sRNA libraries were sequenced by single end (50 bp). The resulting reads were adapter trimmed and filtered on quality by Cutadapt v2.8 [81] and Trimmomatic v0.32 [82], respectively. Additionally, only reads in the 20-25 length range and without indeterminations were kept using a custom script. To study the correlation exhibited by the sRNA expression profiles among the different samples, principal component analysis (PCA) was used. PCA was performed using the prcomp function with scaling in the stats R package v. 4.0.4 (R Core Team 2013). Mann-Whitney-Wilcoxon tests were performed to assess for significant differences in the data clusters for Euclidean distances calculated between groups and among groups with the wilcox.test function in the stats R package. Libraries were aligned to the genomes of HSVd (Y09352.1) and *Cucumis sativus* Chinese Long V3 [83] to determine the provenance of every sRNA. Only sequences that aligned with no mismatches exclusively either to the viroid or the plant genome with a minimum of 0.5 RPMs were considered for further analysis. Additionally, for estimating the size abundance, only samples with all the sRNA lengths were considered. Moreover, HSVd-sRNAs were classified according to their origin (positive or negative viroid strand) considering unique sequences and also the accumulation (rpm). Then, an exact binomial test was performed to assess the significance. Differential expression of cucumber sRNAs was estimated using two R packages DESeq2 [84] and edgeR [85] for pairwise differential expression analysis of expression data. Differentially expressed sRNAs were filtered out using two criteria: (i) adjusted p-value ≤ 0.05 and (ii) RPMs ≥ 5 for at least two libraries in stressed or control samples. P-values were adjusted by False Discovery Rate (FDR). miRNAs were annotated from previously described miRNAs in cucumber [86] and validated with the precursors deposited in RNA central [87]. The rest of sRNA categories were annotated using RNA central and sRNAanno [88].

### mRNA sequencing and analysis

RNA sequencing paired reads (150 bp) were aligned to the cucumber genome using STAR [89]. For non-annotated transcripts, a *de novo* assembly was performed using Trinity [90]. 15589 transcripts non-overlapping with the gene annotation of Chinese Long V3 [83] were aligned with gmap allowing chimeric alignments [91] and a custom annotation of non-coding transcripts was obtained using bedtools [92]. HTSeq-counts [93] was firstly used to count reads per annotated genes using 10 as a minimum alignment quality and secondly of the novel annotated transcripts with a minimum alignment quality of 0 and counting all reads ambiguously mapped in both categories and wherever they overlap (in order to account for potential transposable elements). The count tables were used in DESeq2 [84] to infer significant expression, considering an adjusted p-value under 0.05 for significance. Clustering analysis of the transcripts with significant differential expression in the three time-points was calculated with stats R-package. Firstly, the optimal number of clusters was determined using the complete method in the NbClust R-package considering Euclidean distances [94]. Secondly, the function ‘*kmeans’* of stats was applied with one million iterations. The resulting clusters (grouping transcripts with similar evolution of the differential expression profiles over the infection) were generated by ggplot2 [95]. Moreover, to account for differential exon usage, the package DEXSeq was employed [96], also considering an adjusted p-value under 0.05 for significance. Volcano plots were created using ggplot2 [95]. GO term enrichment analysis was performed using the tool from Cucurbit Genomics Database (CuGenDB) [83]. p-values were corrected using false discovery rate (FDR) and the cutoff p-value for significant represented terms was 0.05.

### Bisulfite sequencing analysis and DMR identification

Whole Genome Bisulfite Sequencing (WGBS) libraries were sequenced by paired end (150 bp). The resulting reads were adapter trimmed and filtered on quality by Trimgalore v0.6.6. Additionally, 10 bases from 5’ ends were trimmed from reads using the same software. The sequence quality was checked by FastQC v0.11.9. The mean conversion rate for the 36 libraries was above 99%. Clean reads were mapped to the reference genome using bismark [98] allowing one mismatch per 25 nucleotides length seed. Alignments at the same position were removed using deduplicate_bismark script, including forward and reverse reads. The bismark_methylation_extractor script was used to extract the methylation call for every single cytosine analyzed and obtain a genome-wide methylation report discriminating by context (CG/CHG/CHH).

The analysis of differentially methylated regions (DMR) was carried out with the R-package DMRcaller [99] dividing the genome in equal bins of 50 bp and pooling the samples of the same condition. The DMR were then computed by performing Fisher’s exact test between the number of methylated reads and the total number of reads in both conditions for each bin. The obtained p-values were then adjusted for multiple testing using Benjamini-Hochberg’s method to control the false discovery [100]. The criteria chosen to consider a bin as a DMR were the following: i) adjusted p-value <= 0.05, ii) at least three cytosines in the specified context, iii) more than a 15% methylation difference between the two conditions and iv) at least an average number of reads of eight. Finally bins that were at less than 300 bp were joined.

In addition to the DMRs identification, overall methylation was calculated as the mean of the percentage methylation of cytosines distinguishing among contexts. Those cytosines covered by less than eight reads were discarded. Next, a heat-map representing the genome-wide methylation difference between the two conditions was performed. First, cytosine methylation percentages were grouped in 3000 nucleotide size windows. Second, the methylation difference was calculated according to time and context. Third, the methylation difference of windows meeting the following terms was set to zero: i) mock or HSVd show zero reads coverage and ii) mock and HSVd show less than eight reads coverage. Methylation difference values less than -15% or greater than 15% were considered as -15% and 15%, respectively. Red and blue colors represented the hypomethylation and hypermethylation in HSVd. Both approaches were also carried out for the ribosomal sequence.

## SUPPLEMENTARY MATERIAL

**Supplementary Figure 1:** Graphic representation of the experimental approach used in this study.

**Supplementary Figure 3:** Viroid-derived sRNAs arise from both HSVd-RNA strands.

**Supplementary Figure 4:** The relative accumulation of 21 and 24 nt vd-sRNAs evolves differentially during the infection.

**Supplementary Figure 5:** Genome view of the vd-sRNAs recovered from infected cucumber plants.

**Supplementary Figure 6:** Two HSVd regions show extremely low accumulation of vd-sRNAs.

**Supplementary Figure 7:** Details of the pipeline used to identify non-annotated differentially expressed cucumber transcripts.

**Supplementary Figure 8:** Gene ontology analysis (plant GOSlim) for transcripts with differential exon usage identified in HSVd-infected plants at 10, 17 and 24 dpi.

**Supplementary Figure 9:** Global view of the temporal evolution of the differential methylation in ribosomal genes in CHG and CHH contexts.

**Supplementary Figure 10:** Graphic representation of the mean coverage of WGBS assay.

**Supplementary Figure 11:** Association between HSVd accumulation and rDNA methylation.

**Supplementary Figure 12:** Association between transcriptional alterations and epigenetic changes induced by HSVd-infection.

**Supplementary Figure 13:** HSVd-infection is associated to significant transcriptional alterations of rRNA.

**Supplementary Table S1:** Description of the oligonucleotides used in this work.

**Supplementary Table S2:** Description of the reads obtained in each analyzed library

**Supplementary Table S3:** Identification and classification of long RNAs-reads.

**Supplementary Table S4:** Identification and classification of the small RNA-reads.

**Supplementary Table S5:** Detailed description of the miRNA family members with significant differential expression in infected plants.

**Supplementary Table S6:** Detailed list of the previously annotated cucumber transcripts with significant differential expression in response to infection.

**Supplementary Table S7:** Detailed list of the previously annotated cucumber transcripts with common significant differential expression in response to infection.

**Supplementary Table S8:** Detailed list of the non-previously annotated cucumber transcripts with significant differential expression in response to infection.

**Supplementary Table S9:** Detailed list of cucumber transcripts with differential exon usage in response to HSVd-infection.

**Supplementary Table S10:** Detailed description of the cucumber gene with at least one DMRs in response to HSVd-infection.

**Supplementary** Table S11: Detailed description of the cucumber genes with at least one antagonistic DMR.

## AUTHOR CONTRIBUTIONS

J.M.M. performed the experiments. P.V.B, J.C.S. and J.M.M performed the bioinformatic analysis. J.M.M., V.P and G.G.: designed the experiment. J.M.M., P.V.B, J.C.S., V.P. and G.G. analyzed and discussed the results. G.G. Conceived the general experiment and drafted the manuscript. All authors read, revise and approved the final manuscript.

## ACKNOWLEDGMENTS

The authors thank Dr. Beatriz Navarro for critically reading this manuscript.

## DATA AVAILABILITY

All data needed to evaluate the conclusions in the paper are present in the paper and/or the Supplementary Materials. Additional data related to this paper may be requested from the corresponding author. All sequence datasets have been uploaded in NCBI SRA under the accession code BioProject ID: PRJNA795883

