## Supplementary Figures for "Multiomic analisys reveals that viroid infection induces a temporal reprograming of plant-defence mechanisms at multiple regulatory levels"

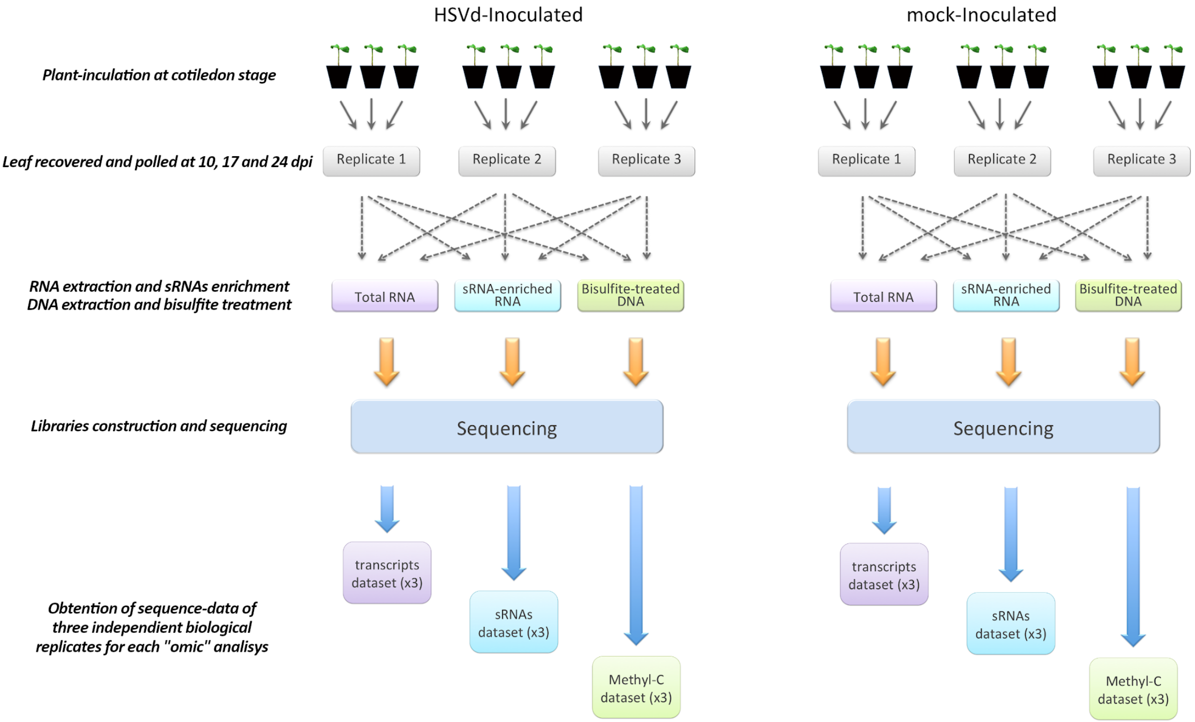


**Figure S1:** Graphic representation of the experimental approach used in this study to identify HSVd-induced alterations in the cucumber transcriptome, sRNAnome and methylome. In total 54 independent libraries corresponding to three biological replicates for HSVd- and mock-inoculated samples at three different time points (10 dpi, 17 dpi and 24 dpi) were analyzed.

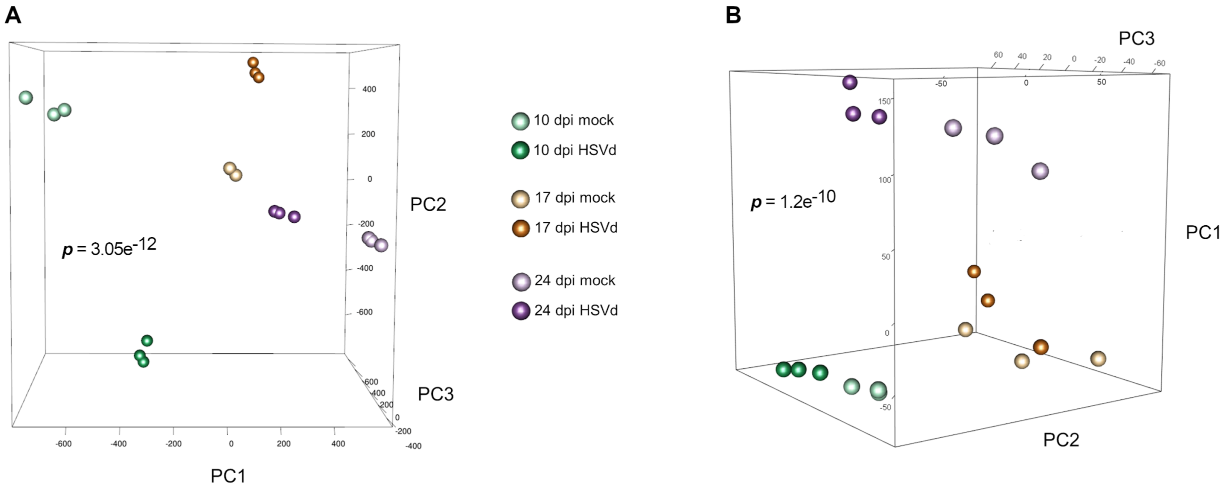


**Figure S2:** Principal component analysis based on sRNAs (A) and long transcripts (B) accumulation in biological replicates of HSVd-infected and mock-inoculated plants at the three analyzed time-points. The percentages of variance explained by the first three principal components (PC) were 16.43%, 15.01% and 14.47%, respectively in A (45,91% of the total variance) and 40.25%, 14.37% and 7.9% in B (62.52% of th etotal variance). The statistical significance was estimated by Mann-Whitney-Wilcoxon test, considering the inter- and intra-group Euclidean distances.

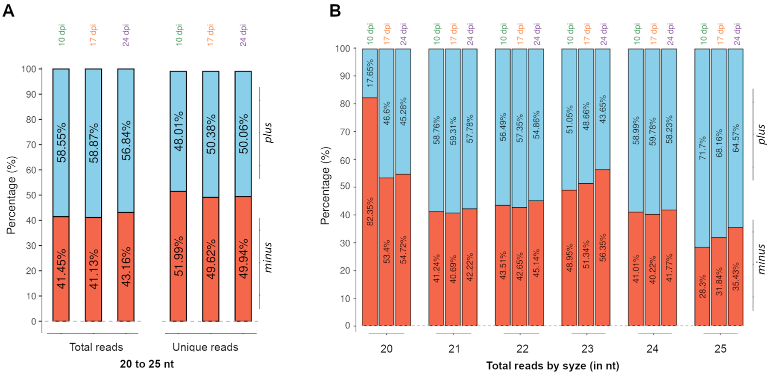


**Figure S3: Viroid-derived sRNAs arise from both HSVd-RNA strands. A**) Relative accumulation of the total (left) and unique (right) vd-sRNAs (20 to 25 nt) derived from genomic (*plus*) and antigenomic (*minus*) HSVd-RNA strands at the three analyzed time points. **B**) Polarity distribution of the total vd-sRNAs discriminated by infection time and size.

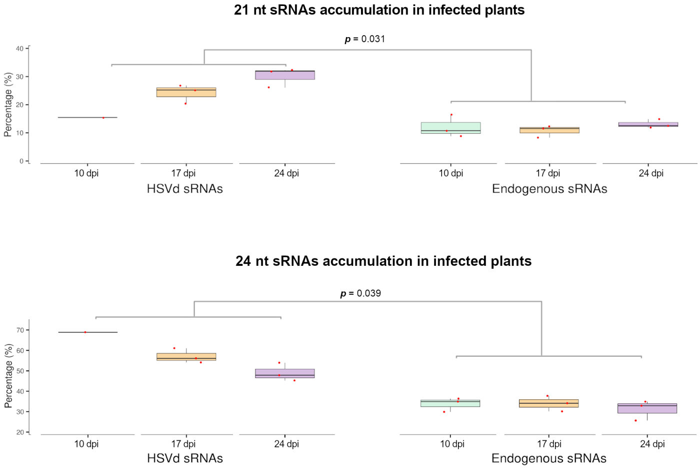


**Figure S4: The relative accumulation of 21 and 24 nt vd-sRNAs evolves differentially during the infection**. Box-plot analysis showing a comparison of the mean values (in percentage) observed for 21 nt (upper panel) and 24 nt (lower panel) vd-sRNAs with endogenous sRNAs of the corresponding size at 10, 17 and 24 dpi. The expression of each sRNA type is represented by the median (internal box line) of the percentage of RPM values. The significance of the differences was estimated by a 2-ways non-parametric ANCOVA test.

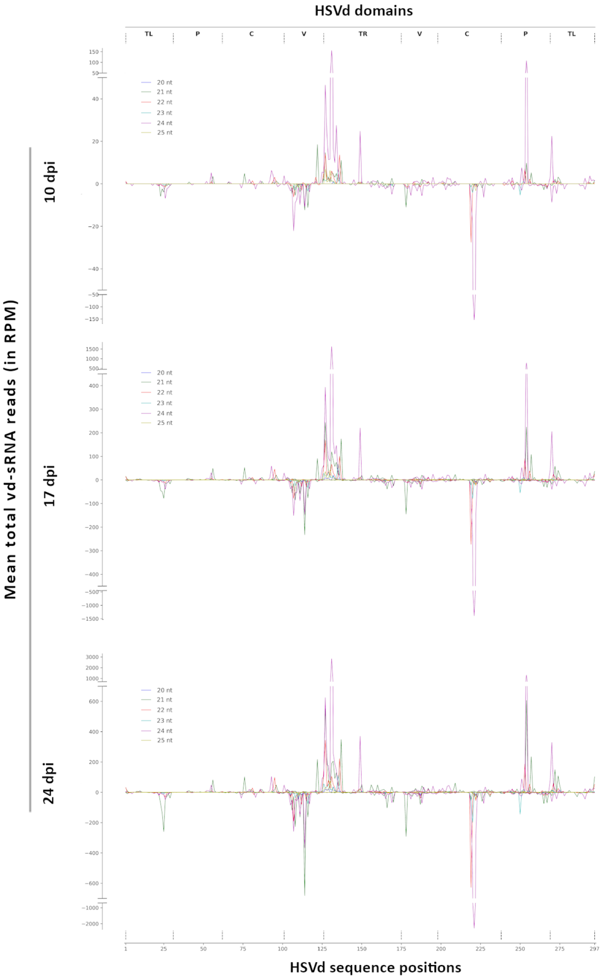


**Figure S5: Genome view of the vd-sRNAs recovered from infected cucumber plants.** The vd-sRNAs were plotted according to the position of their 5´-end onto the HSVd- RNA sequence in either sense (above the x-axis) or antisense (below the x-axis) configuration. The values on the y-axis represent the mean of total vd-sRNAs (sizes indicated by colors) in the three analyzed time points.

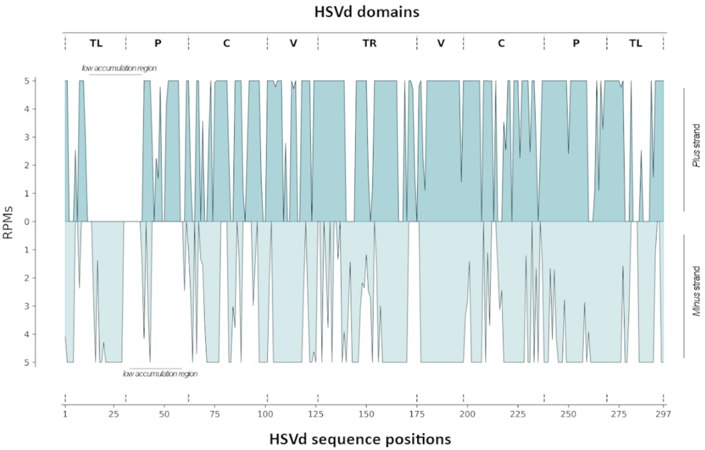


**Figure S6: Two HSVd regions show extremely low accumualtion of vd-sRNAs.** Detailed genome view of the vd-sRNAs recovered from infected cucumber plants. The vd-sRNAs (21 to 25 nt) were plotted according to the position of their 5´-end onto the HSVd- RNA sequence in either sense (above the x-axis) or antisense (below the x-axis) configuration. The values on the y-axis represent the abundance of total vd-sRNAs limited to 5 RPM in order to evaluate which regions of the HSVd genome contribute to the production of vd-sRNAs.

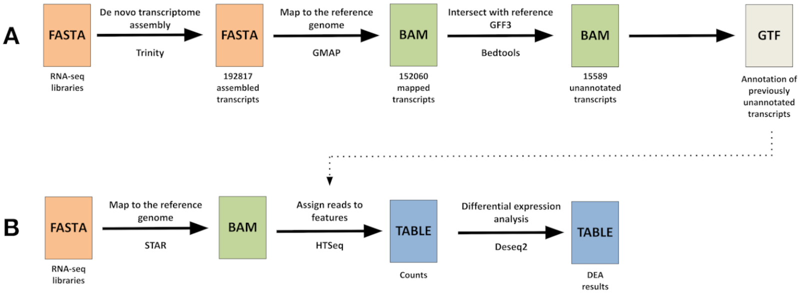


**Figure S7: Details of the pipeline used to identify non-annotated differentially expresed cucumber transcripts. A**) Assembly and annotation of previously non-identified transcripts. **B**) Alignment of reads to cucumber genome and differential expression analysis.

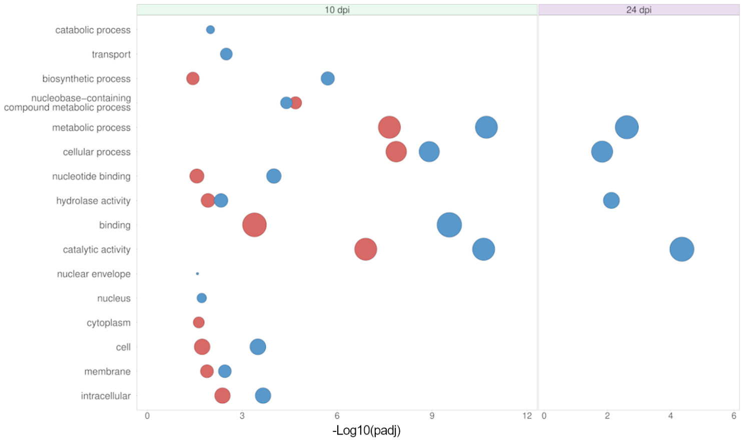


**Figure S8: Gene ontology analysis (plant GOSlim) for transcripts with differential exon usage identified in HSVd-infected plants at 10, 17 and 24 dpi.** Circle size represents level of enrichment and color down- (red) or up-regulated (blue) response. The -Log(10) of the adjusted P values is represented in the X-axis.

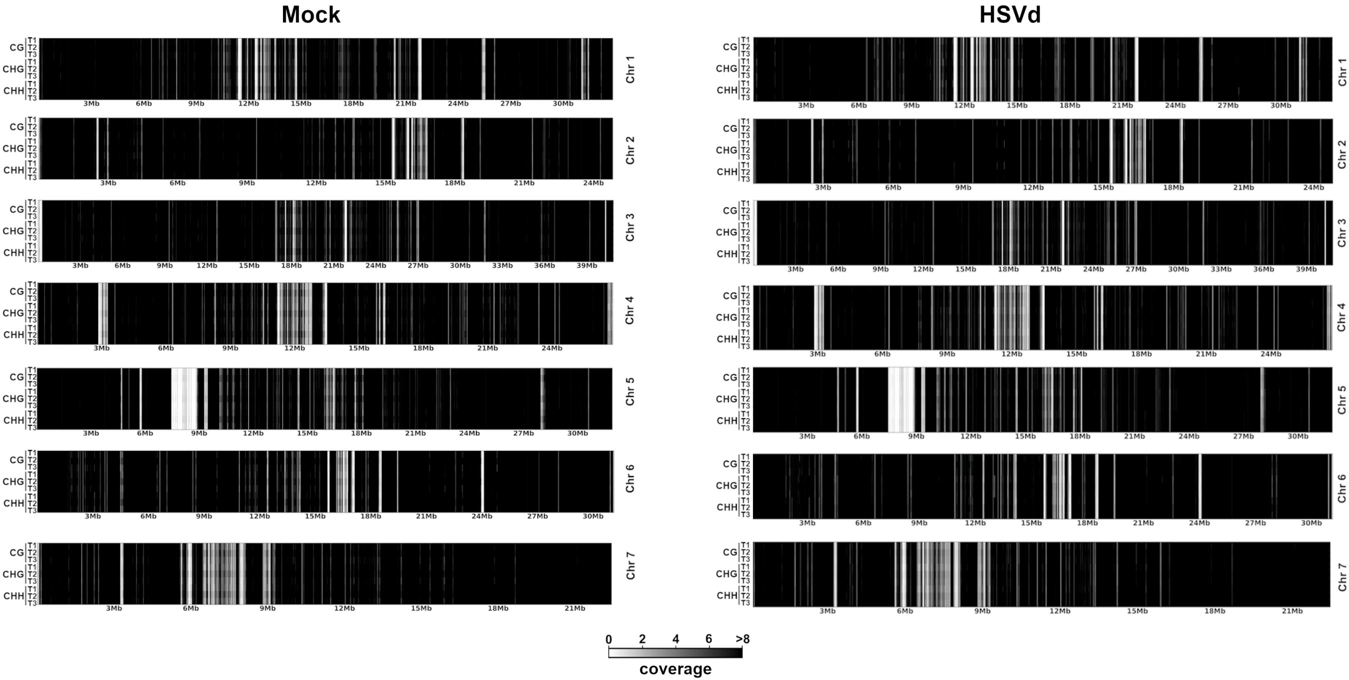


**Supplementary Figure 9**. Graphic representation of the mean coverage by chromosome of WGBS sequencing for mock (left) or HSVd (right) samples. The mean coverage for each methylation context (CG,CHG and CHH) is plotted at the three time points, being T1 = 10 dpi, T2 = 17 dpi and T3 = 24 dpi. The coverage is represented by the mean number of reads of each position in 3 kb windows.

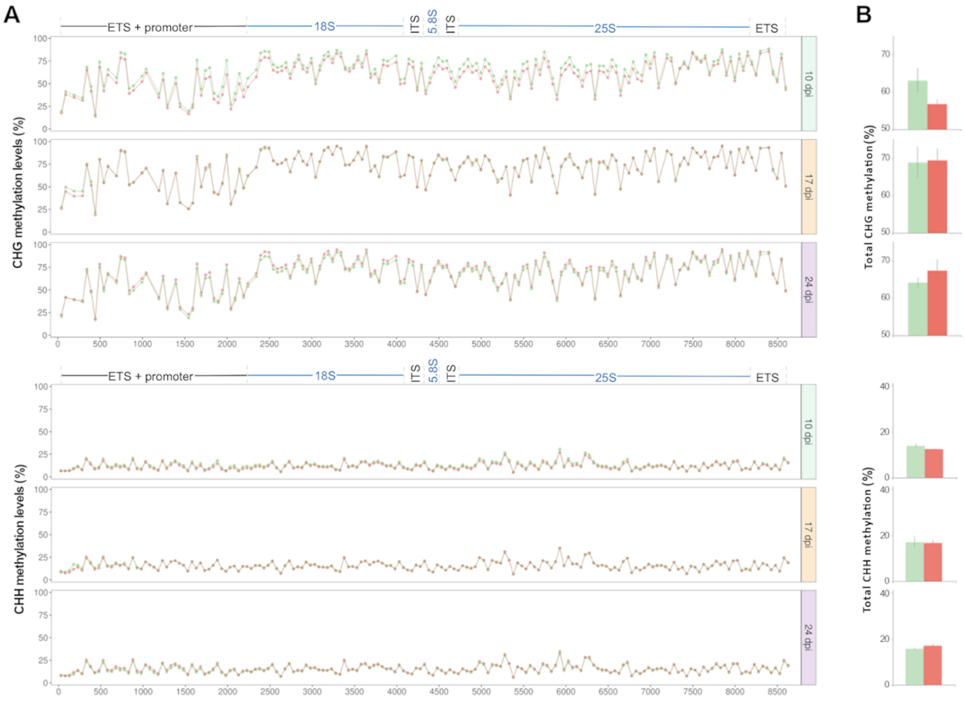


**Figure S10: Global view of the temporal evolution of the differential methylation in ribosomal genes in CHG and CHH contexts. A**) Methylation profiles in CHG (upper) and CHH (lower) context associated to HSVd-infection. Red and green dots represent the means of methylation (estimated for 50 nt regions) of rDNA in HSVd-infected and control plants respectively. B) Graphic representation of the means of the total cytosine methylation values at CHG (upper) and CHH (lower) sequence context of ribosomal genes in HSVd-infected and control plants at 10, 17 and 24 dpi. The ribosomal genes (18S, 5.8S and 25S) are indicated in blue while the internal transcribed spacers (ITS) and external transcribed spacers (ETS) are in black.

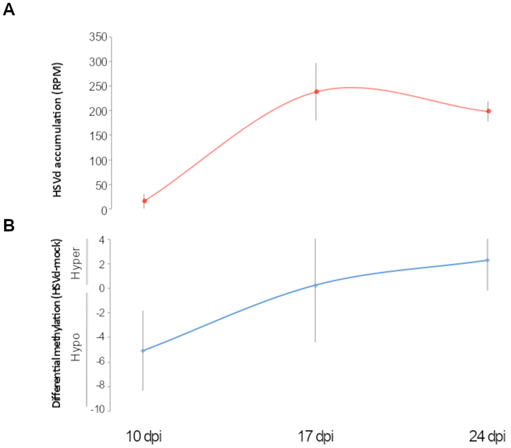


**Figure S11: Association between HSVd accumulation and rDNA methylation.** **A**) Representation of the temporal evolution of viroid accumulation estimated by the mean of HSVd-transcript reads recovered in each analyzed time (in red). **B**) Graphic showing the mean differential methylation in infected plants of ribosomal genes at the three analyzed time points. Error bars indicate the standard error between replicates.

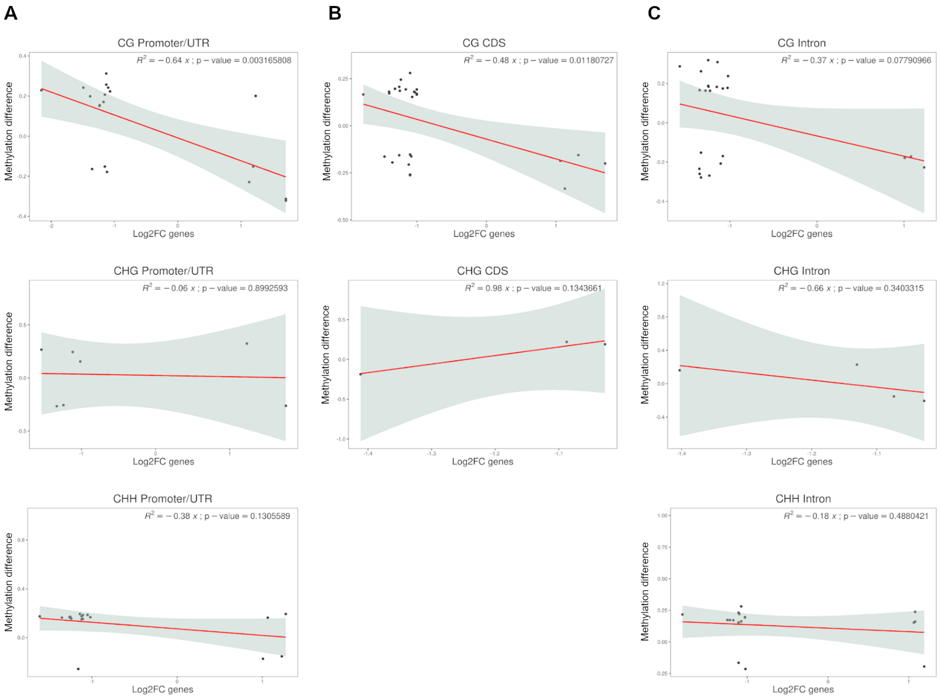


**Figure S12: Association between transcriptional alterations and epigenetic changes induced by HSVd-infection**. Scatter plots showing the correlation (with signifcance estimated by *Pearson correlation coefficient*) between the expression levels of differential genes containing at least one antagonistic DMR and their global methylation considering the three selected gene regions (**A**) Promoter/UTR, (**B**) CDS and (**C**) intron in the three sequence contexts: CG (upper), CHG (middle) and CHH (lower).

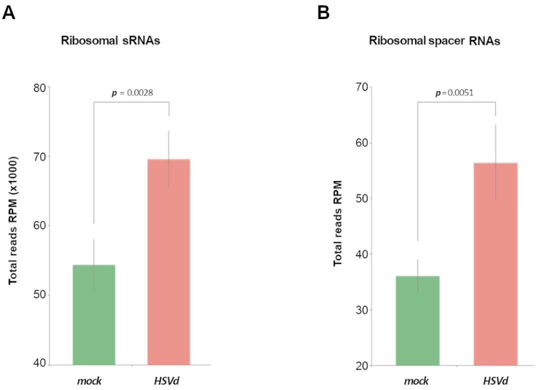


**Figure S13: HSVd-infection is associated to significant transcriptional alterations of rRNA**. Diagrams showing the mean accumulation of the total ribosomal derived sRNAs (**A**) and transcripts of the ribosomal internal spacers (**B**) recovered from mock-incoculated (green) and HSVd-infected (red) plants during the analyzed infection period. Error bars indicate the SE between biological replicates. Significance values were estimated by paired T-test.
